# Dysregulation of gene expression during gastrulation results in impaired primitive erythropoiesis and vascular development in Trim71-KO embryos

**DOI:** 10.1101/2024.07.12.603214

**Authors:** Tobias Beckröge, Bettina Jux, Hannah Seifert, Hannah Theobald, Elena De Domenico, Stefan Paulusch, Marc Beyer, Andreas Schlitzer, Elvira Mass, Waldemar Kolanus

**Affiliations:** Molecular Immunology and Cell Biology, Life and Medical Sciences Institute (LIMES), University of Bonn, Bonn, Germany; Quantitative Systems Biology, Life & Medical Sciences Institute (LIMES), University of Bonn, Bonn, Germany; Genomics and Immunoregulation, Life & Medical Sciences Institute (LIMES), University of Bonn, Bonn, Germany; Systems Medicine, Deutsches Zentrum für Neurodegenerative Erkrankungen (DZNE) e.V., Bonn, Germany; PRECISE Platform for Genomics and Epigenomics, Deutsches Zentrum für Neurodegenerative Erkrankungen (DZNE) e.V. and University of Bonn and West German Genome Center, Bonn, Germany; Immunogenomics & Neurodegeneration, Deutsches Zentrum für Neurodegenerative Erkrankungen (DZNE) e.V., Bonn, Germany; Developmental Biology of the Immune System, Life & Medical Sciences Institute (LIMES), University of Bonn, Bonn, Germany

## Abstract

The transition of an embryo from gastrulation to organogenesis requires precisely coordinated changes in gene expression. The RNA-binding protein Trim71 is essential for embryonic survival, but its exact role in mammalian development *in vivo* remains poorly defined. Here we show that murine *Trim71*-KO embryos appear normal until embryonic day (E)8.5 but display severe defects in primitive erythropoiesis, yolk sac vasculature and heart function during the onset of organogenesis at E9.5 and E10.5. This led to an impaired vascular translocation of yolk sac-derived macrophage progenitors to the embryo head, independent of Trim71 expression in erythro-myeloid progenitors. The cardiovascular and erythropoiesis defects explain the embryonic lethality upon global *Trim71*-KO. Targeting *Trim71* in hematoendothelial progenitors did not induce strong developmental defects, indicating an earlier developmental origin of these phenotypes in *Trim71*-KO embryos. ScRNA-seq of E7.5 *Trim71*-KO embryos revealed that transcriptomic changes arise already at gastrulation, showing a strong upregulation of the transcription factor Eomes. We identify Eomes as a direct target of Trim71-mediated mRNA repression via the NHL domain, demonstrating a functional link of Trim71 to a key regulator of mesodermal development. Taken together, our data suggest that Trim71-dependent control of gene expression at gastrulation establishes a framework for proper development during organogenesis.

## Introduction

The circulatory system consists of blood vessels, the heart and blood cells, and it mediates the exchange of oxygen, nutrients and cells throughout the body. The components of the circulatory system develop at the onset of organogenesis, starting at E8.5 in mice (Bautch and Caron 2015).

Cardiovascular development and the generation of blood cells from hematopoiesis are essential for mammalian embryonic survival during organogenesis (Shalaby et al. 1995; Dumont et al. 1994; Koushik et al. 2001; Fujiwara et al. 1996). Cells of the circulatory system are derived from the mesodermal germ layer, which emerges from cells passing through the primitive streak during gastrulation at E6.5–E7.5 in murine embryonic development (Prummel et al. 2020; Bardot and Hadjantonakis 2020). Mesodermal progenitor cells that leave the primitive streak give rise to hematoendothelial progenitors (HEP), which can differentiate into endothelial cells (EC) that form the inner lining of blood vessels (Pijuan-Sala et al. 2019). The extraembryonic yolk sac generates a vast vascular network through the assembly of EC into a primitive capillary plexus (vasculogenesis, E7.5–E8.5) and subsequent remodeling of the vascular network by vessel sprouting, pruning and increase in vessel diameter (angiogenesis, from E8.5 on) (Garcia and Larina 2014). By primitive erythropoiesis, HEP can also directly give rise to primitive erythroid cells (EryP) that carry oxygen and remain the only erythroid cell population until E11.5 in mice (Ema et al. 2006; Pijuan-Sala et al. 2019; Lux et al. 2008; Iturri et al. 2021; McGrath and Palis 2008). EryP are first present in the yolk sac and disseminate through the embryo body in parallel to the establishment of blood circulation following the onset of heart function (McGrath et al. 2003). Transient definitive hematopoietic cells also emerge in the yolk sac by the differentiation of EC into erythro-myeloid progenitors (EMP) at E8.5 (McGrath et al. 2015; Kasaai et al. 2017). EMP give rise to pre-macrophages (pMac), that exit the yolk sac via the vasculature and translocate into the embryo proper, where they give rise to intraembryonic macrophage populations (Mass et al. 2016; Stremmel et al. 2018).

Recent studies have highlighted the importance of RNA-binding proteins in cardiovascular development (Völkers et al. 2024). The RNA-binding protein Trim71 is an essential and conserved regulator of embryonic development (Mitschka et al. 2015; Ecsedi et al. 2015; Lin et al. 2007). At the molecular level, Trim71 controls post-transcriptional gene expression by the interaction of its NHL domain with secondary structures in the 3ˈ UTR of mRNAs that have been termed Trim71 responsive elements (TREs) (Kumari et al. 2018; Torres-Fernández et al. 2019). Target mRNA binding by Trim71 leads to their degradation, thus repressing gene expression (Loedige et al. 2013). We and others have previously reported that global knockout of *Trim71* (*Trim71*-KO) in mice results in embryonic lethality at E9.5–E11.5 (Mitschka et al. 2015; Maller Schulman et al. 2008; Cuevas et al. 2015). *Trim71*-KO embryos display a cranial neural tube closure defect, but it has been argued before that this is presumably not the cause of lethality (Ecsedi and Grosshans 2013). The underlying reason for the embryonic lethality upon *Trim71*-KO is so far not understood. While the molecular functions of Trim71 have been extensively studied *in vitro* (Worringer et al. 2014; Welte et al. 2023), the role of Trim71 in mammalian embryonic development *in vivo* is largely unexplored. So far, a role of mammalian Trim71 has been described in neurogenesis and germ line development (Chen et al. 2012; Torres-Fernández et al. 2021; Du et al. 2020). Mutations in the human *TRIM71* gene cause congenital hydrocephalus, highlighting the relevance of this gene for human prenatal development (Duy et al. 2022; Duy et al. 2024). Considering the widespread expression of *Trim71* at gastrulation and early organogenesis (Chen et al. 2012), it is plausible that Trim71 also has functions in the development of other cell types, beyond neural and germ cells, that have not yet been described.

In the present study, we identify Trim71 as an essential factor for primitive erythropoiesis and cardiovascular development, explaining the embryonic lethality of murine *Trim71*-KO embryos. Surprisingly, expression of *Trim71* in HEP and EC is largely dispensable for cardiovascular development and EryP generation. Instead, we show that *Trim71*-KO results in extensive transcriptional changes in the mesoderm at E7.5 that directly precede the onset of defects in the hematoendothelial cell lineage. Mechanistically, Trim71 antagonizes the expression of the mesodermal pioneer transcription factor Eomes and binds to Eomes mRNA in an NHL-domain dependent manner, indicating Trim71-mediated post-transcriptional repression of *Eomes*. Our results delineate novel functions of Trim71 in the mesoderm and indicate that defects in the development of the circulatory system can be initiated at gastrulation.

## Results

### Impaired primitive erythropoiesis and vascular development in *Trim71*-KO embryos

To characterize the onset and progression of developmental phenotypes caused by global deficiency of Trim71, the morphology of *Trim71*-KO embryos was evaluated by light microscopy at E7.5–E10.5. *Trim71*-KO embryos were morphologically indistinguishable from wildtype littermate control (WT) embryos at E7.5 and E8.5 (Fig. 1A). The visible formation of a primitive streak at E7.5, a hallmark of gastrulation, was not impeded by deletion of *Trim71* (Fig. 1A). *Trim71*-KO embryos were normal in size and appearance at E8.5 and acquired the characteristic embryonic morphology until E9.5 (Fig. 1A), indicating normal ventral folding morphogenesis and axial rotation (Gavrilov and Lacy 2013). In agreement with previous observations, *Trim71*-KO embryos were smaller than WT embryos at E9.5, and displayed a cranial neural tube closure defect (Fig. 1A) (Mitschka et al. 2015). These phenotypes were even more pronounced at E10.5, at which stage *Trim71*-KO embryos showed severe underdevelopment across the whole body (Fig. 1A). The growth retardations after E9.5 were underscored by decreased cell numbers in the yolk sac, embryo head and embryo body (Supplemental Fig. S1A–C). In addition to these previously described phenotypes, we observed that *Trim71*-KO embryos appear pale in color from E9.5 on. This was in particular noticeable in the heart and the dorsal aorta, which are normally red in color in WT embryos due to the presence of EryP (Fig. 1A). This prompted us to further analyze the yolk sac of *Trim71*-KO embryos, which is the origin of all erythroid cells before E11.5 (McGrath and Palis 2008). Likewise, *Trim71*-KO yolk sacs also appeared pale (Fig. 1B) and a quantification of relative EryP numbers by flow cytometry at E9.5 and E10.5 revealed a strong reduction of EryP in *Trim71*-KO compared to WT yolk sacs (Fig. 1C,D). A significant decrease of EryP numbers was also observed in the embryo head and embryo body at E10.5 (Fig. 1E,F). Altogether, these data show that *Trim71*-KO leads to a defect in primitive erythropoiesis.

**Figure 1:**
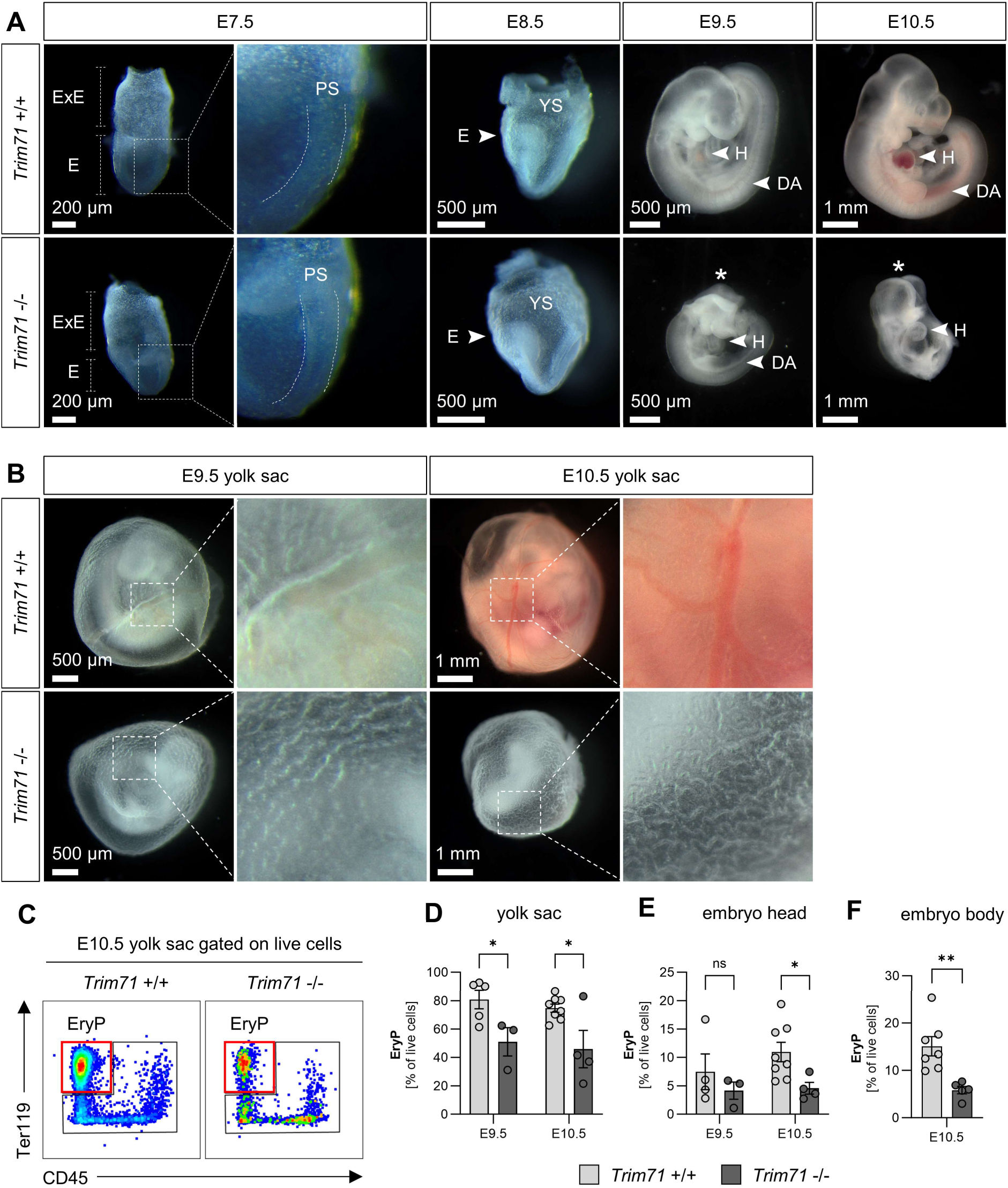
*Trim71*-KO embryos show developmental retardation and decreased primitive erythropoiesis at early organogenesis. (A) Morphology of WT and *Trim71*-KO embryos at indicated developmental stages E7.5–E10.5. Dashed boxes show magnification of the primitive streak region at E7.5. E = embryo, ExE = Extraembryonic region, DA = dorsal aorta, H = heart, PS = primitive streak, YS = yolk sac. Stars indicate the presence of a neural tube closure defect. (B) Morphology of WT and *Trim71*-KO yolk sacs at E9.5 and E10.5. Dashed boxes show magnification of vitelline vessels. (C) Representative flow cytometry gating of Ter119^+^ CD45^-^ EryP in E10.5 WT and *Trim71*-KO yolk sacs. Red boxes indicate gates for EryP. (D–F) Relative quantification of EryP in the (D) yolk sac and (E) embryo head at E9.5 and E10.5, and (F) the embryo body at E10.5 by flow cytometry (n = 3–8 embryos from 2–3 experiments, data depicted as mean±SEM, unpaired Student’s *t*-test, (*) *P* < 0.05, (**) *P* < 0.01).

The yolk sac is an early site of vascular development and is an integral component of the embryonic circulatory system. After E9.5, the yolk sac vasculature is composed of large blood vessels, also known as vitelline vessels, and microvascular areas (Garcia and Larina 2014). Closer inspection of the light microscopy images of E9.5 and E10.5 *Trim71*-KO yolk sacs revealed abnormal vascular structures (Fig. 1B). We further investigated the yolk sac vasculature by whole mount immunofluorescence staining using the EC marker CD31. Overview microscopy images showed that *Trim71*-KO yolk sacs were completely devoid of large vitelline blood vessels and had instead only equally-sized small blood vessels (Fig. 2A). Moreover, higher-magnification imaging of the yolk sac microvasculature showed clear structural differences in vascular network upon *Trim71*-KO (Fig. 2B). We quantified endothelial extensions and vascular branching points in the microvasculature as markers of yolk sac angiogenesis. Endothelial extensions, defined as CD31^+^ structures emerging from one vessel but not yet connected to another vessel (Fig. 2B magnification), were strongly reduced upon *Trim71*-KO (Fig 2C). These endothelial extensions can represent newly sprouting vessels or regressing vessels, two key angiogenic processes (Jones et al. 2008). Likewise, *Trim71*-KO led to a significant reduction in branching points, defined as the intersection of at least three vessels (Fig. 2B magnification, Fig. 2D). The relative numbers of EC in the yolk sac, embryo head and embryo body were however not affected by *Trim71*-KO (Supplemental Fig. S1D–F). Taken together, these data demonstrate that *Trim71-*KO embryos have a yolk sac remodeling defect, whereas vasculogenesis per se appears to be intact. We further investigated if the intraembryonic circulatory system is affected by *Trim71*-KO. *Trim71*-KO embryos form a heart (Fig. 1A), but 50% of E9.5 and 20% of E10.5 *Trim71*-KO embryos had no heartbeat (Fig. 2E, Supplemental Movie 1, 2). Furthermore, the subset of E10.5 *Trim71*-KO embryos that had a heartbeat showed a significant reduction in heart rate compared to WT embryos (Fig. 2F, Supplemental Movie 3). In summary, *Trim71*-KO embryos display defects in all major components of the circulatory system.

**Figure 2:**
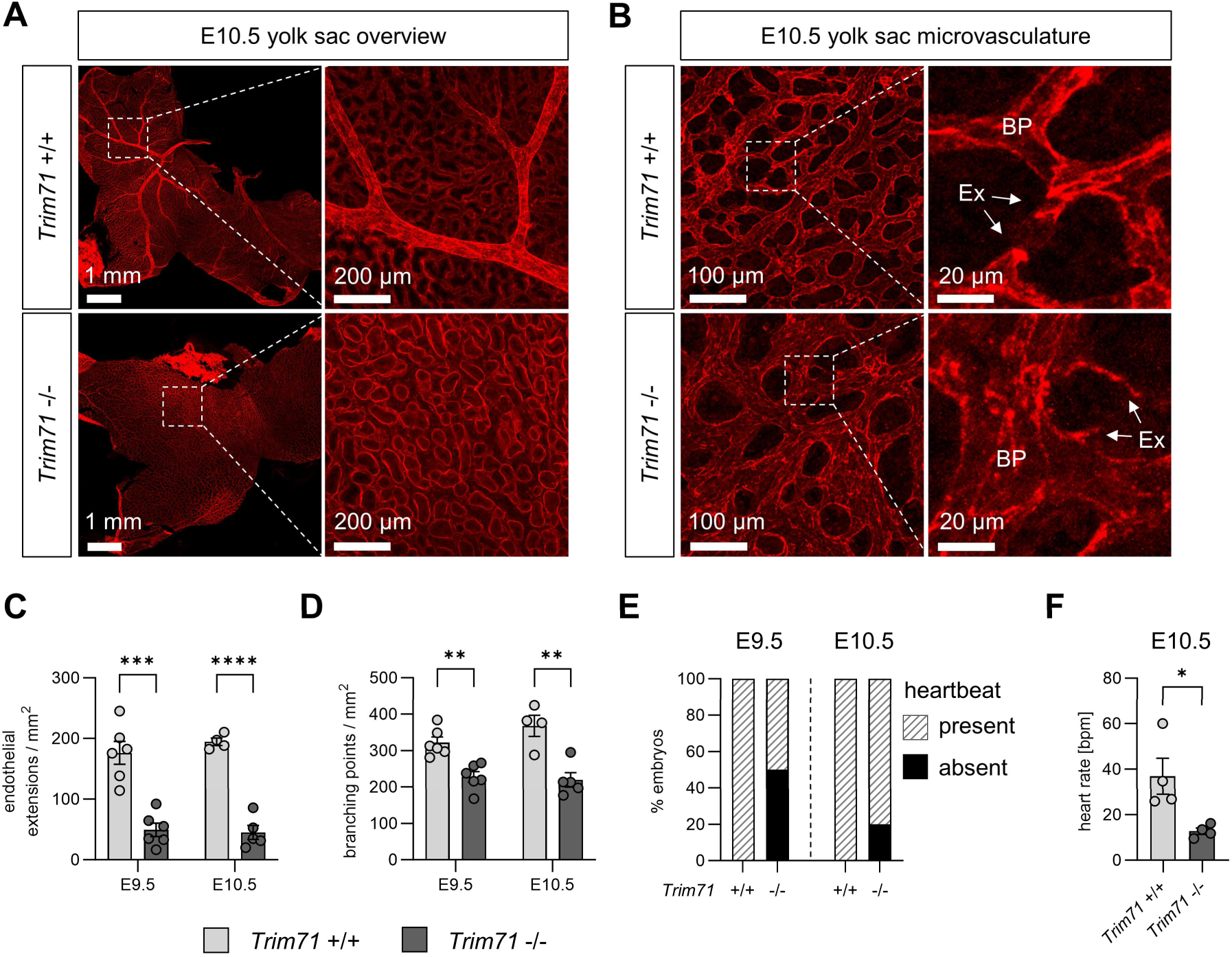
Cardiovascular defects in *Trim71*-KO embryos. (A) Representative overview images of E10.5 yolk sacs stained with the EC marker CD31. Dashed boxes show magnification of vitelline vessels in *Trim71*-WT yolk sac, whereas a representative region devoid of vitelline vessels is shown in the *Trim71*-KO yolk sac. (B) Representative images of the yolk sac microvasculature at E10.5 stained with CD31. Dashed boxes show magnification of individual vessels with indicated endothelial extensions and branching points. BP = branching point, Ex = endothelial extension. (C,D) Quantification of (C) endothelial extensions and (D) branching points in the yolk sac microvasculature at E9.5 and E10.5 (n = 4–6 yolk sacs from 3 experiments per stage, data depicted as mean±SEM, unpaired Student’s *t*-test, (**) *P* < 0.01, (***) *P* < 0.001), (****) *P* < 0.0001). (E) Percentage of embryos with present or absent heartbeat at E9.5 and E10.5 (n = 8–25 embryos from 5–14 experiments). (F) Heart rate of E10.5 WT and *Trim71*-KO embryos in which a heartbeat was detectable (n = 4 embryos from 3 experiments, data depicted as mean±SEM, unpaired Student’s *t*-test, (*) *P* < 0.05).

### Defects in the circulatory system result in impaired vascular pMac translocation in *Trim71*-KO embryos

In order to investigate the functional relevance of the circulatory system defects in *Trim71*-KO embryos, we analyzed the translocation efficiency of EMP-derived pMac from the yolk sac to the embryo proper, where they give rise to intraembryonic macrophage populations (Mass et al. 2016). The translocation of pMac occurs through the yolk sac vasculature and depends on proper heart function (Stremmel et al. 2018; Ginhoux et al. 2010). Definitive hematopoietic cell numbers in the yolk sac, embryo body and embryo head were analyzed by flow cytometry. In *Trim71*-KO yolk sacs, EMP numbers were slightly increased at E9.5 and pMac numbers were elevated at E10.5, whereas yolk sac macrophage numbers were unaffected (Fig. 3A). In contrast, pMac and macrophages were markedly reduced in the body and were almost completely absent from the head of *Trim71*-KO embryos (Fig. 3B,C). These data indicate that transient definitive hematopoietic cells emerge normally in *Trim71*-KO yolk sacs, but pMac fail to translocate from the yolk sac to the embryo proper. Besides cardiovascular defects, impaired macrophage progenitor translocation can also result from hematopoietic cell-intrinsic defects, for example caused by the loss of the chemokine receptor Cx3cr1 (Mass et al. 2016). To address this possibility, we first analyzed *Trim71* expression in transient definitive hematopoietic cells of WT embryos. Indeed, we found that *Trim71* is expressed by EMP and pMac, while its expression declines upon differentiation towards macrophages (Fig. 3D). We therefore tested if targeted deletion of *Trim71* in EMP and their progeny, using the *Csf1r*^iCre^ driver line and the *Trim71*-flox line, has an effect on EMP-derived hematopoiesis or intraembryonic macrophage colonization. *Csf1r*^iCre^ *Trim71* conditional knockout (cKO) embryos did not show any differences in the number of EMP, pMac and macrophages in the yolk sac and embryo head at E9.5 and E10.5 compared to control embryos (Fig. 3E,F). Moreover, *Csf1r*^iCre^ *Trim71* cKO pMac and macrophages showed no changes in the expression of Cx3cr1 (Supplemental Fig. S2A–C). These data demonstrate that expression of *Trim71* in EMP is not required for intraembryonic macrophage colonization, and rule out an EMP- or pMac-intrinsic origin of the impaired vascular translocation of pMac from the yolk sac into the embryo proper upon *Trim71*-KO.

**Figure 3:**
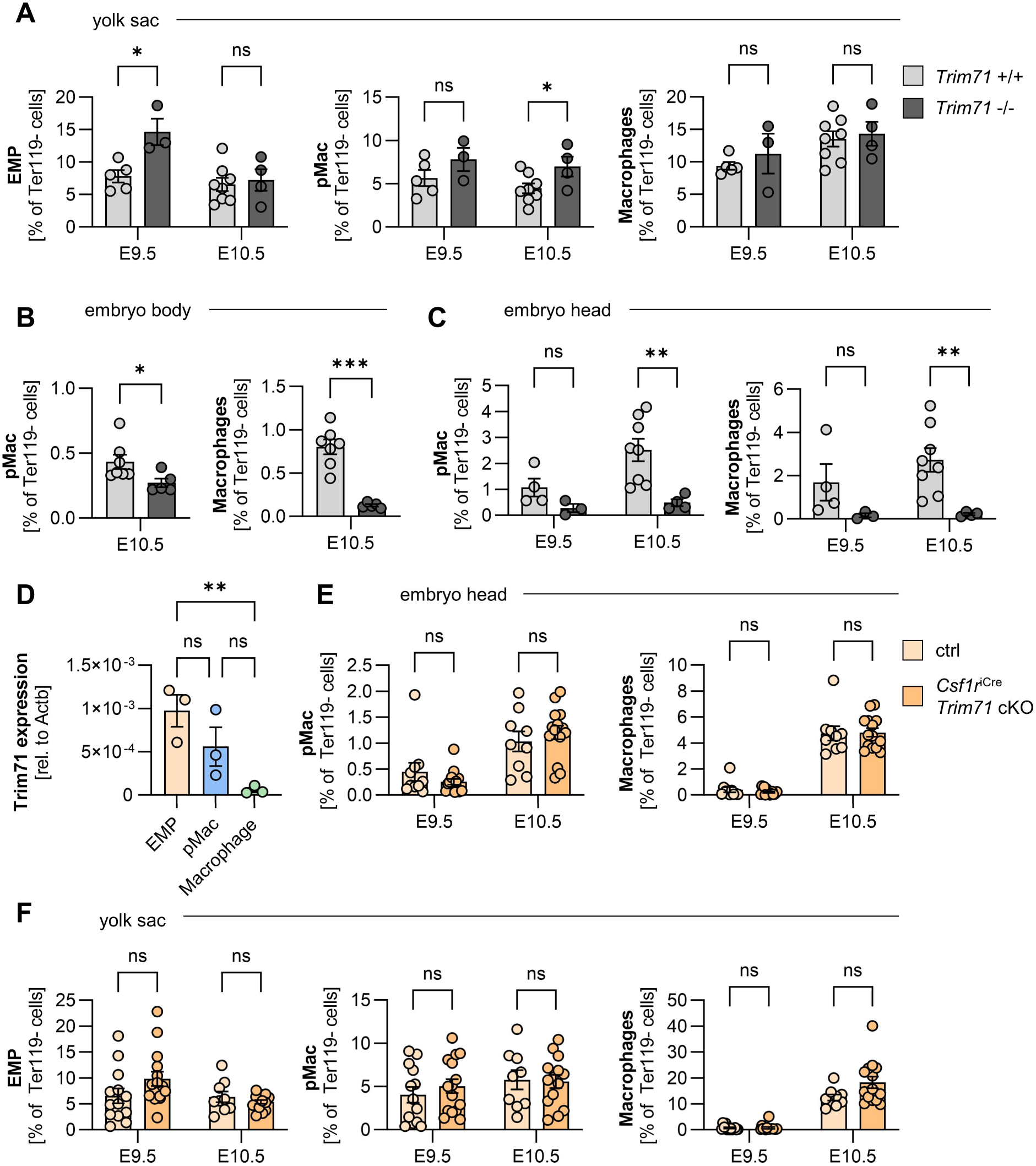
Effect of global or EMP-specific *Trim71* deletion on EMP-derived myeloid cell numbers in the yolk sac and the embryo. (A–C) Quantification of hematopoietic cell numbers in WT and *Trim71*-KO embryos. (A) Relative numbers of EMP, pMac and macrophages in the yolk sac at E9.5 and E10.5. (B) Relative numbers of pMac and macrophages in the embryo body at E10.5 and (C) in the embryo head at E9.5 and E10.5 (n = 3–8 embryos from 2–3 experiments, data depicted as mean±SEM, unpaired Student’s *t*-test, ns = not significant, (*) *P* < 0.05, (**) *P* < 0.01). (D) Quantification of Trim71 mRNA expression by qRT-PCR in EMP, pMac and macrophages isolated from WT E10.5 yolk sacs (n = 3 from 3 experiments, data depicted as mean±SEM, ordinary one-way ANOVA, ns = not significant, (**) *P* < 0.01). (E,F) Quantification of hematopoietic cell numbers in control and *Csf1r*^iCre^ *Trim71* cKO (*Csf1r*^iCre/+^ *Trim71*^fl/fl^) embryos. Ctrl indicates *Csf1r*^+/+^ *Trim71*^fl/fl^. (E) Relative numbers of pMac and macrophages in the embryo head at E9.5 and E10.5. (F) Relative numbers of EMP, pMac and macrophages in the yolk sac at E9.5 and E10.5 (n = 9–15 embryos from 4–5 experiments, data depicted as mean±SEM, unpaired Student’s *t*-test, ns = not significant).

### Yolk sac scRNA-seq identifies transcriptional changes in *Trim71*-KO EC

To analyze changes in gene expression associated with impaired vascular development of *Trim71*-KO embryos, we performed single-cell mRNA-sequencing (scRNA-seq) of whole E9.5 WT and *Trim71*-KO yolk sacs. We identified the cell types expected to be present in the yolk sac, including EC (Fig. 4A, Supplemental Fig. 3A). Uniform manifold approximation and projection (UMAP) showed differential distribution of cells from the same cell type between genotypes, indicating widespread transcriptomic changes in *Trim71*-KO yolk sacs (Fig. 4B). EC showed extensive changes in gene expression, with 430 downregulated differentially expressed genes (DEG) upon *Trim71*-KO (Supplemental Fig. 3B). Fitting to the flow cytometry data (Fig. 1C), EryP were proportionally decreased upon *Trim71*-KO in the scRNA-seq dataset, while no changes in EC numbers were present (Supplemental Fig. 3C,D). Gene ontology (GO) analysis of downregulated DEG in EC revealed an enrichment of genes in multiple processes related to angiogenesis (Fig. 4C). Downregulated genes contained in the process *regulation of angiogenesis* included the endothelial transcription factor Ets1, which is known to play a role in vascular development (Wei et al. 2009), and the blood-flow induced transcription factor Klf2 (Lee et al. 2006) (Fig. 4D). Moreover, *Trim71*-KO EC had decreased expression of the cell junction protein Cdh5 (Fig. 4D), which is required for proper angiogenesis and can be transactivated by Ets1 (Bentley et al. 2014; Lelièvre et al. 2000). In line with the decreased endothelial extensions in *Trim71*-KO yolk sacs (Fig. 2C), the gene expression score of genes involved in s*prouting angiogenesis* was significantly decreased in *Trim71*-KO EC (Fig. 4E). Visualization of GO-processes from downregulated genes in EC by a category network plot showed the presence of four clusters, which were related to cell migration, endothelial cell differentiation, cell junction assembly and regulation of vascular development (Fig. 4F). These data indicate that the impaired yolk sac vascular remodeling of *Trim71*-KO embryos is the result of multiple distinct EC-intrinsic processes involved in angiogenesis.

**Figure 4:**
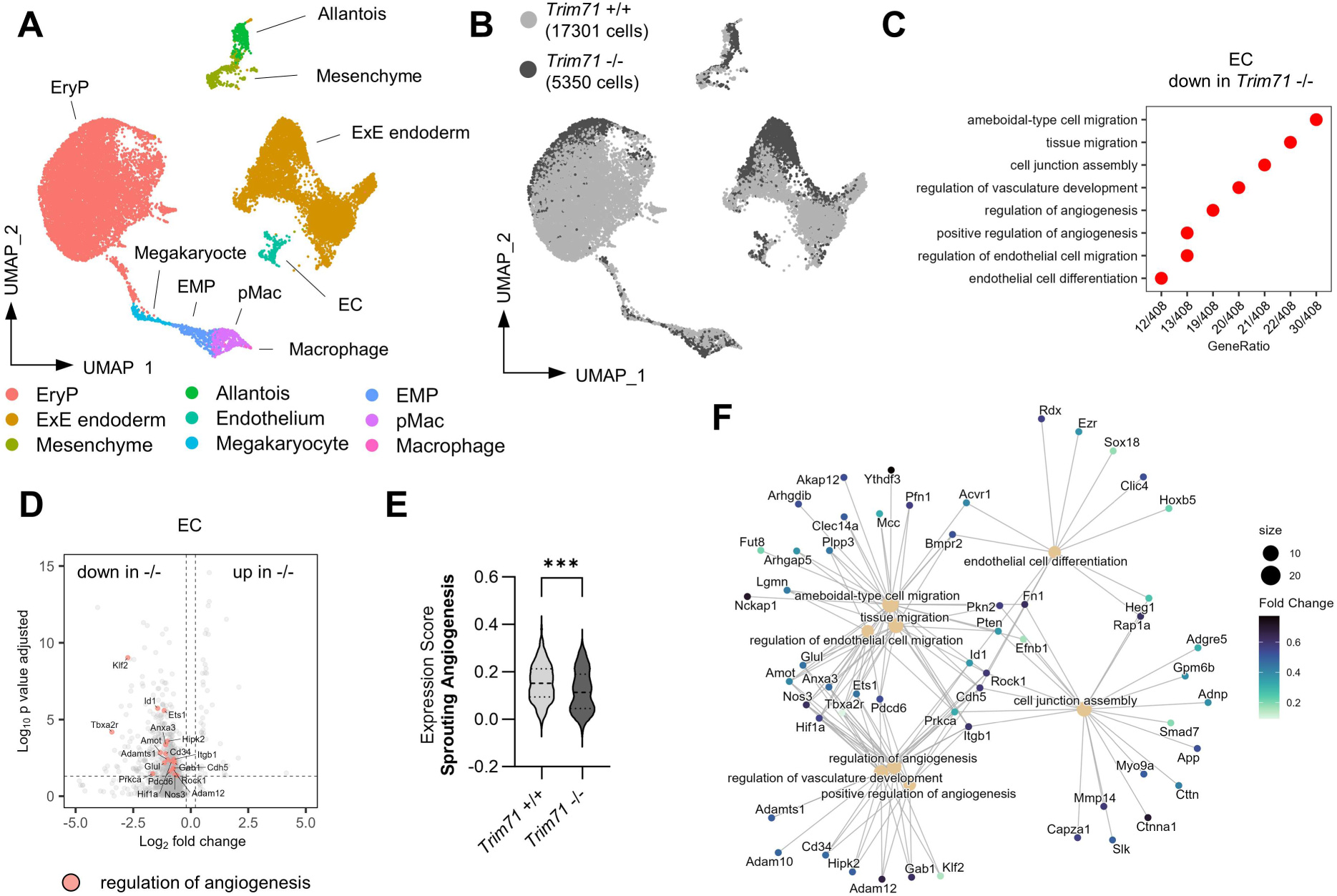
ScRNA-seq of E9.5 yolk sacs reveals decreased endothelial expression of angiogenic genes upon *Trim71*-KO. (A) UMAP plot of all cells with color-coded cell type annotations. (B) UMAP plot with color-coded genotypes (light gray = *Trim71*^+/+^, dark gray = *Trim71*^-/-^). (C) Enriched GO-terms among downregulated DEG in *Trim71*^-/-^ EC. (D) Volcano plot of DEG in *Trim71*^-/-^ EC with genes included in the GO-term *regulation of angiogenesis* highlighted in red. (E) Expression score of the MSigDB gene set *sprouting angiogenesis* (data depicted as violin plot with median and quartiles indicated as dashed lines, unpaired Student’s t-test, (***) *P* < 0.001). (F) Category network plot of GO-terms from DEG downregulated in *Trim71*^-/-^ EC with color-coded expression fold changes of genes included in the processes. The circle size denotes the number of DEG contained in each process.

### Trim71 expression in the mesoderm at gastrulation and in the hematoendothelial lineage

We next sought to investigate the developmental origins leading to the impaired vascular development and defective primitive erythropoiesis of *Trim71*-KO embryos. To this end, we examined Trim71 expression from gastrulation to early organogenesis with a focus on the hematoendothelial cell lineage. Analysis of scRNA-seq data from E6.5–E8.5 WT mouse embryos (Pijuan-Sala et al. 2019) by pseudo-bulk expression analysis across cell types showed that *Trim71* is highly expressed in the primitive streak and nascent mesoderm (Supplemental Fig. 4A). Intermediate *Trim71* expression levels were present in HEP and EC, whereas *Trim71* expression was absent in EryP (Supplemental Fig. 4A). Using immunofluorescence staining, we confirmed the protein expression of Trim71 in Flk1^+^ cells of E7.5 WT embryos, which encompass both mesodermal progenitors and HEP (Biben et al. 2023) (Supplemental Fig. 4B). Moreover, Trim71 protein expression was also present in CD31^+^ embryonic EC of the yolk sac at E9.5 and the dorsal aorta at E10.5 (Supplemental Fig. 4C,D).

### *Tie2*^Cre^ induced *Trim71* deletion does not phenocopy *Trim71*-KO

Since we detected the expression of Trim71 in HEP and EC, we investigated the function of Trim71 in these cells by conditional knockout induced via the *Tie2*^Cre^ driver line (*Tie2*^Cre^ *Trim71* cKO). Tie2 is expressed in HEP at late gastrulation (Ema et al. 2006), and the *Tie2*^Cre^ line targets both EC and EryP (Kisanuki et al. 2001; Tang et al. 2010). At E10.5, *Tie2*^Cre^ *Trim71* cKO embryos and yolk sacs appeared normal in overall morphology (Fig. 5A). Mendelian ratios of *Tie2*^Cre^ *Trim71*^fl/fl^ were retrieved at developmental stages E9.5–E12.5 and viable *Tie2*^Cre^ *Trim71*^fl/fl^ were born, showing that *Tie2*^Cre^ *Trim71* cKO does not lead to embryonic lethality (Fig. 5B). In the yolk sac, *Tie2*^Cre^ *Trim71* cKO did not prevent the formation of large vitelline vessels (Fig. 5A,C). We detected a slight decrease in endothelial extensions and branching points in the yolk sac microvasculature upon *Tie2*^Cre^ *Trim71* cKO at E9.5 (Fig. 5D,E), these effects were however mild compared to global *Trim71*-KO (Fig. 2B–D). At E12.5, endothelial extensions remained significantly reduced in *Tie2*^Cre^ *Trim71* cKO yolk sacs, whereas branching points normalized to control embryo levels (Fig. 5D,E). All *Tie2*^Cre^ *Trim71* cKO embryos had a heartbeat at E9.5 and E10.5 (Fig. 5F) and showed no changes in relative EryP numbers in the yolk sac (Fig. 5G). These results demonstrate that Trim71 expression in HEP is dispensable for primitive erythropoiesis and heart function, and is only marginally required for yolk sac angiogenesis.

**Figure 5:**
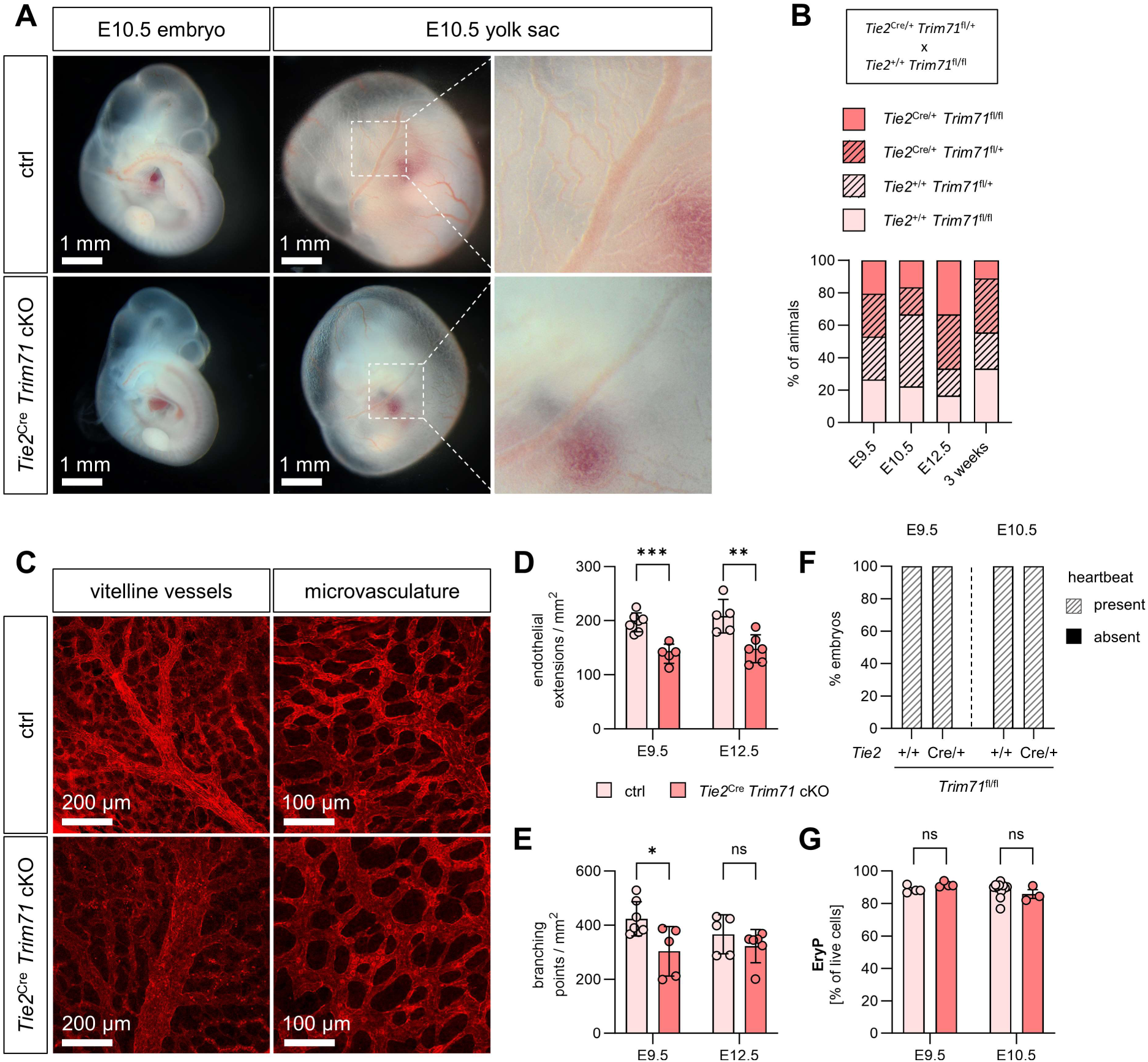
*Tie2*^Cre^ *Trim71* cKO does not result in embryonic lethality and does not induce strong defects in cardiovascular development or primitive erythropoiesis. (A) Morphology of control or *Tie2*^Cre^ *Trim71* cKO (*Tie2*^Cre/+^ *Trim71*^fl/fl^) embryos and yolk sacs at E10.5. Ctrl indicates *Tie2*^+/+^ *Trim71*^fl/fl^ or *Tie2*^+/+^ *Trim71*^fl/+^. Dashed boxes show magnification of vitelline vessels. (B) Quantification of genotype percentages in E9.5, E10.5 and E12.5 embryos and 3-week-old pups from mating of *Tie2*^Cre/+^ *Trim71*^fl/+^ with *Tie2*^+/+^ *Trim71*^fl/fl^ mice (n = 9–34 embryos or pups from 2–5 litters). (C–E) Whole mount staining of yolk sacs with CD31. (C) Representative images of vitelline vessels and microvascular areas of E9.5 yolk sacs. Quantification of (D) endothelial extensions and (E) branching points in the microvasculature of E9.5 and E12.5 yolk sacs (n = 5–6 yolk sacs from 3–5 experiments, data depicted as mean±SEM, unpaired Student’s *t*-test, ns = not significant, (*) *P* < 0.05, (**) *P* < 0.01, (***) *P* < 0.001). (F) Percentage of embryos with present or absent heartbeat at E9.5 and E10.5 (n = 3–12 embryos from 2–4 experiments). (G) Relative quantification of EryP in the yolk sac at E9.5 and E10.5 by flow cytometry (n = 3– 11 embryos from 1–2 experiments, data depicted as mean±SEM, unpaired Student’s *t*-test, ns = not significant).

### Transcriptomic alterations in *Trim71*-KO embryos arise at gastrulation

The absence of strong vascular and erythropoiesis phenotypes in *Tie2*^Cre^ *Trim71* cKO embryos led us to the hypothesis that the origin of the defects observed in *Trim71*-KO embryos could lie earlier in embryonic development during gastrulation. We thus performed scRNA-seq of whole E7.5 *Trim71*-KO embryos. Cell type annotation led to the identification of the expected gastrulation stage cell types, including cells from all three germ layers (Fig. 6A, Supplemental Fig. 5A). *Trim71*-KO embryos had normal numbers of cells from all germ layers, demonstrating successful gastrulation (Supplemental Fig. 5B,C). Moreover, in WT embryo cells Trim71 was highly and ubiquitously expressed across all germ layers (Supplemental Fig. 5D). We focused on the mesoderm for differential gene expression analysis, since the hematoendothelial cell lineage is derived from this germ layer. Remarkably, *Trim71*-KO E7.5 mesodermal progenitors already displayed extensive transcriptomic changes, as evident by a slightly different relative position of WT and *Trim71*-KO mesodermal cells on the UMAP plot (Fig. 6B) and a substantial amount of DEG (Fig. 6C, 29 upregulated DEG, 67 downregulated DEG). GO-analysis of downregulated genes in *Trim71*-KO mesoderm yielded GO-terms related to mRNA processing, RNA splicing, RNA stability and chromatin remodeling (Fig. 6D–F). Furthermore, we analyzed upregulated genes in *Trim71*-KO mesoderm in order to identify mRNAs that are directly targeted for degradation by Trim71 during gastrulation. *Trim71*-KO resulted in elevated mesodermal expression of the transcription factors Eomes and Lhx1, both of which play important roles in mesodermal development and are contained in the GO-term *mesendoderm development* (Fig. 6F–I). Increased expression of Eomes was not only restricted to mesodermal cells, but was also present in cells of the primitive streak and endodermal cells (Fig. 6H). Lhx1 was not expressed in primitive streak cells in either cell type, but was increased upon *Trim71*-KO in mesodermal and endodermal cells (Fig. 6I). These data show that changes in mesodermal gene expression at late gastrulation precede the onset of defects in the circulatory system of *Trim71*-KO embryos.

**Figure 6:**
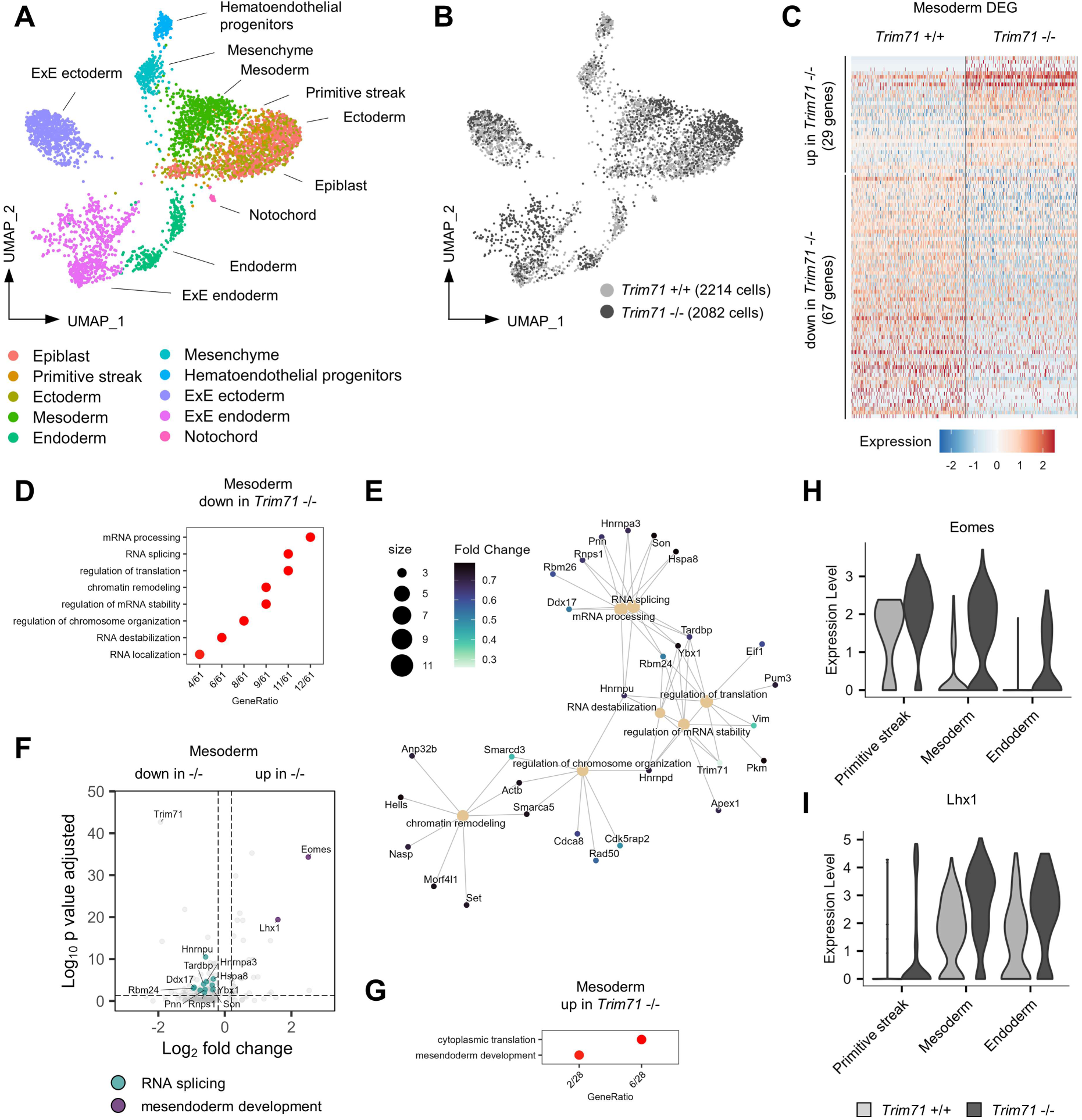
ScRNA-seq of E7.5 embryos reveals extensive transcriptional changes in the mesoderm upon *Trim71*-KO. (A) UMAP plot of all cells with color-coded cell type annotations. (B) UMAP plot with color-coded genotypes (light gray = *Trim71*^+/+^, dark gray = *Trim71*^-/-^). (C) Expression heatmap of upregulated and downregulated DEG in *Trim71*^+/+^and *Trim71*^-/-^ mesodermal cells. (D) Enriched GO-terms among downregulated DEG in *Trim71*^-/-^ mesodermal cells. (E) Category network plot of GO-terms from DEG downregulated in *Trim71*^-/-^ mesodermal cells with color-coded expression fold changes of genes included in the processes. The circle size denotes the number of DEG contained in each process. (F) Volcano plot of DEG in *Trim71*^-/-^ mesodermal cells with DEG included in the GO-term *RNA splicing* highlighted in turquoise and *mesendodermal development* in purple. (G) Enriched GO-terms among upregulated DEG in *Trim71*^-/-^ mesodermal cells. (H,I) Expression of (H) Eomes and (I) Lhx1 in cells of the primitive streak, mesoderm and endoderm separated by genotype.

### Trim71 antagonizes Eomes expression and binds to Eomes mRNA dependent on the NHL domain

We used mESC as an *in vitro* model to study a potential direct regulation of Eomes and Lhx1 expression by Trim71. Cells were differentiated within embryoid bodies towards the mesodermal lineage for four days (Pearson et al. 2008). We confirmed increased expression of Eomes in *Trim71*-KO compared to *Trim71*-flox mESC, that express WT levels of Trim71, at day 4 of differentiation (Fig. 7A). In contrast, Lhx1 expression was unaltered between genotypes (Fig. 7B). RNA secondary structure prediction showed the presence of a putative Trim71 responsive element (TRE) within the 3ˈ UTR of the murine Eomes mRNA, that fulfils the structural requirements for interaction with the Trim71 NHL domain (Fig. 7C) (Kumari et al. 2018; Shi et al. 2024; Torres-Fernández et al. 2019). The sequence of this TRE was UAUCUUGGAGAUA, located at position 3452–3465 within the Eomes mRNA (Fig. 7D). In agreement with reported TRE characteristics (Kumari et al. 2018), the Eomes mRNA TRE consisted of a 13-mer stem-loop with a U-A base pair at the top of the stem and a G in position III of the loop (Fig. 7D). To test direct binding of Eomes mRNA by Trim71, we performed cross-linking immunoprecipitation (CLIP) using mESC lines endowed with endogenous expression of mNeon-FLAG-tagged *Trim71*-WT, *Trim71*-KO or *Trim71*-R595H variants (Duy et al. 2022) at day 4 of differentiation (Fig. 7E,F). The *Trim71*-R595H mutation is located within the NHL domain and abrogates the RNA binding capacity of Trim71 (Duy et al. 2022). Flow cytometric analysis of Trim71 expression via the mNeon tag validated the absence of Trim71 expression in *Trim71*-KO mESC, while expression was retained in *Trim71*-WT and *Trim71*-R595H cells (Fig. 7G,H). Strikingly, FLAG-CLIP of Trim71 variants in differentiated mESC showed an enrichment of Eomes mRNA in *Trim71*-WT mESC, which was strongly reduced upon *Trim71*-KO and *Trim71*-R595H mutation (Fig. 7I). Lhx1 mRNA was not reliably detected by qPCR after CLIP (Fig. 7I). Importantly, we validated Trim71 protein enrichment after immunoprecipitation (Fig. 7J). Together, these data show that Trim71 selectively interacts with Eomes mRNA in an NHL-domain dependent manner, suggesting a regulation of this key mesodermal transcription factor by the previously described mechanisms of Trim71-induced mRNA degradation (Loedige et al. 2013; Torres-Fernández et al. 2019).

**Figure 7:**
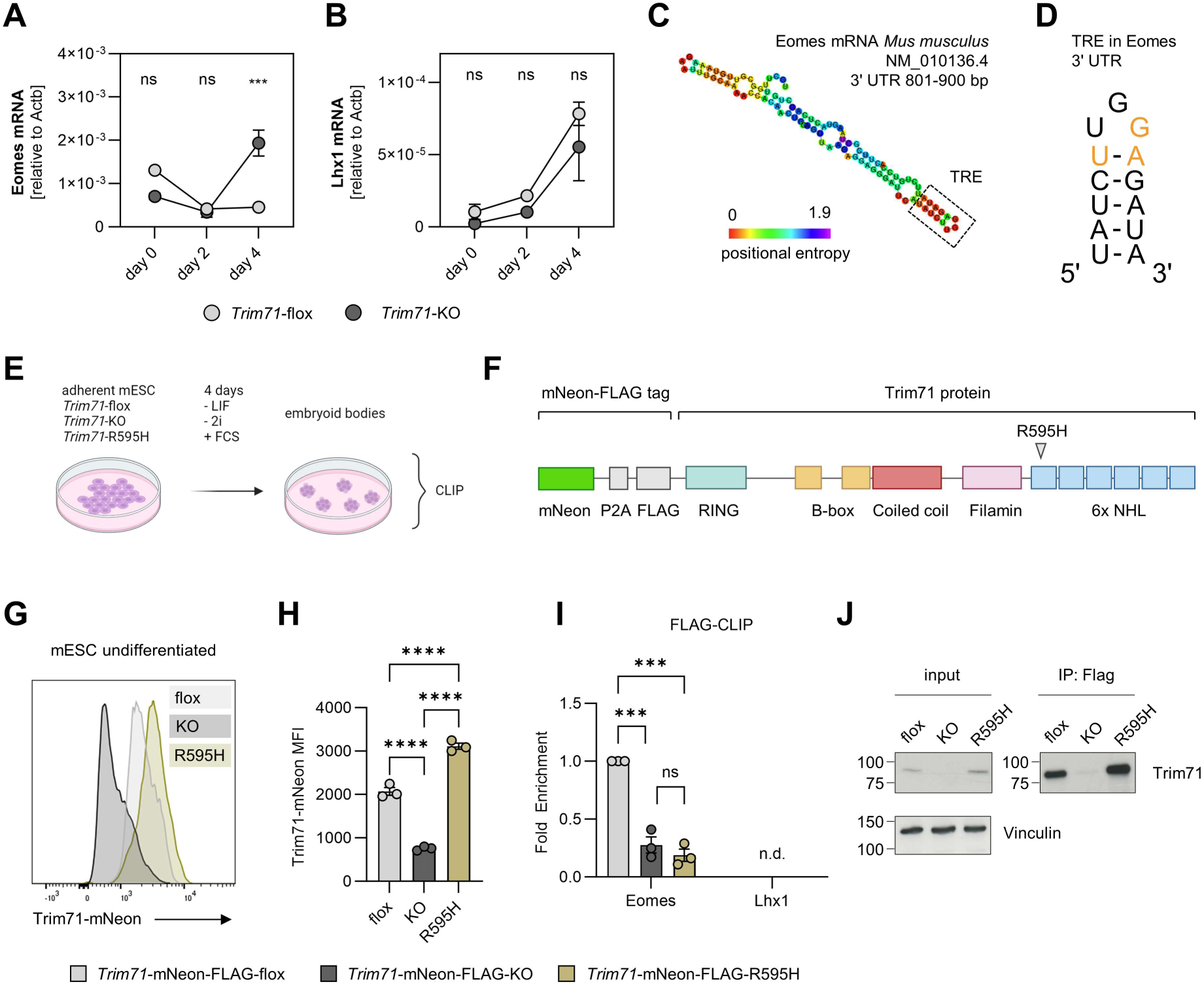
Trim71 antagonizes Eomes expression by binding its mRNA via the NHL domain. (A– B) Expression of (A) Eomes and (B) Lhx1 in *Trim71*-WT and *Trim71*-KO mESC at day 0, day 2 and day 4 of differentiation as determined by qRT-PCR (n = 3, data depicted as mean±SEM, two-way ANOVA, ns = not significant, (***) *P* < 0.001). (C) RNAfold secondary structure prediction of the 801–900 bp region of the murine Eomes mRNA 3ˈ UTR. The identified TRE is highlighted by a dashed box. (D) Representation of the identified TRE in the Eomes mRNA 3ˈ UTR. Critical bases for the interaction with the Trim71 NHL domain are highlighted in orange. (E) Depiction of *Trim71*-mNeon-FLAG-WT, -KO or -R595H mESC differentiation in embryoid bodies, followed by CLIP at day 4. Created with BioRender.com. (F) Representation of the Trim71-mNeon-FLAG protein with indicated domains and position of the R595H mutation. (G) Representative histogram and (H) quantification of the median fluorescence intensity (MFI) of the Trim71-mNeon signal in undifferentiated *Trim71*-mNeon-FLAG-WT, -KO or -R595H mESC (n = 3, data depicted as mean±SEM, ordinary one-way ANOVA, (****) *P* < 0.0001). (I) Quantification of Eomes and Lhx1 fold enrichment in FLAG-CLIP of *Trim71*-mNeon-FLAG-WT, -KO or -R595H mESC at day 4 of differentiation (n = 3, data depicted as mean±SEM, one-way ANOVA, ns = not significant, (***) *P* < 0.001). (J) Western blot of input and immunoprecipitated protein lysates after FLAG-CLIP detected with Trim71 and Vinculin antibodies.

## Discussion

### Trim71 is required for primitive erythropoiesis and cardiovascular development

In this study, we identify Trim71 as a crucial factor for the embryonic development and function of all major components of the circulatory system, including EryP, the vasculature and the heart. In global *Trim71*-KO embryos, defects in the circulatory system become apparent with the onset of organogenesis at E9.5. We show that *Trim71*-KO embryos are noticeably pale and have strongly reduced levels of EryP in the yolk sac and in the embryo proper. Severe defects in vascular development were present in the yolk sac, as evident by the complete absence of large vitelline vessels and an unremodeled vascular network. Vasculogenesis was not noticeably impeded by loss of Trim71, but angiogenesis was severely decreased in *Trim71*-KO yolk sacs, as shown by decreased vascular branching points and a loss of endothelial extensions in the yolk sac microvasculature. These phenotypes are comparable to the yolk sac vascular remodeling defects of embryos deficient for crucial angiogenic factors, such as *Tie2*^-/-^, *Nrp1*^-/-^ or *Notch1*^-/-^ embryos (Tachibana et al. 2005; Jones et al. 2008; Krebs et al. 2000). At the transcriptional level, scRNA-seq of E9.5 yolk sacs revealed decreased endothelial expression of genes involved in cell migration, cell junction assembly and endothelial cell differentiation upon *Trim71*-KO. These processes are collectively required for vascular development, angiogenesis and sprouting (Lamalice et al. 2007; Bazzoni and Dejana 2004), providing an explanation for the loss of endothelial extensions and the yolk sac remodeling defect. Besides vascular impairments, *Trim71*-KO embryos display defects in heart function, as evident from a partial absence of heartbeat and a decreased heart rate. This was associated with defective colonization of the embryo proper by macrophages and their progenitors (pMac), that originate from the yolk sac. The presence of normal pMac and macrophage numbers in the embryo head of *Csf1r*^iCre^ *Trim71* cKO embryos demonstrates that the translocation defect of these cells in *Trim71*-KO embryos is independent of the Trim71 expression in EMP and pMac. This strongly indicates that the impaired heart function of these embryos leads to insufficient blood circulation to mediate the translocation of pMac through the vasculature (Stremmel et al. 2018; Ginhoux et al. 2010), given that the embryonic heart rate correlates with blood flow velocity (Phoon et al. 2000). Since the vitelline vessels are the direct vascular connection between the yolk sac and the embryo, the atrophy of these blood vessels in *Trim71*-KO embryos might further enhance the retention of pMac within the yolk sac. An impaired blood circulation of *Trim71*-KO embryos and the reduced hematocrit due to the absence of EryP could additionally restrict vascular remodeling, which is known to be dependent on the hemodynamic forces of the blood (Garcia and Larina 2014; Koushik et al. 2001; Jones et al. 2008). Concordantly, we observed decreased expression of the blood flow induced transcription factor Klf2 in *Trim71*-KO EC (Lee et al. 2006). Thus, impaired vascular remodeling in *Trim71*-KO embryos is the result of both EC-intrinsic loss of angiogenic activity and the reduction of external hemodynamic forces.

As argued before, the cranial neural tube closure observed in *Trim71*-KO embryos is likely not the cause of their embryonic lethality at E9.5–E11.5, considering that embryos carrying other genetic ablations leading to neural tube defects typically survive until the late fetal period (Copp et al. 2003; Maller Schulman et al. 2008). In contrast, vascular development, primitive erythropoiesis and proper heart function are indispensable for mid-gestational survival of the embryo, and deficiencies in either process lead to lethality at the onset of organogenesis (Coultas et al. 2005; Shalaby et al. 1995; Fujiwara et al. 1996; Koushik et al. 2001). Furthermore, the detection of defects in these processes in *Trim71*-KO embryos temporally coincides with the appearance of developmental retardation and morphological anomalies from E9.5 on. Considering that defects in either component of the circulatory system are sufficient to drive embryonic lethality, the combined defects in all major parts of the circulatory system of *Trim71*-KO embryos provides an adequate explanation for their developmental arrest and lethality.

### Trim71-dependent control of gene expression during gastrulation determines proper vascular development and primitive erythropoiesis

We further investigated the developmental origin of the vascular and erythropoiesis defects in *Trim71*-KO embryos. Trim71 was expression was present in mesodermal progenitors, HEP and EC, but was absent in EryP. Using *Tie2*^Cre^ *Trim71* cKO embryos, we show that expression of Trim71 in HEP and their progeny is dispensable for embryonic survival, heart function and primitive erythropoiesis, and only marginally affects yolk sac angiogenesis. We therefore considered the mesoderm during gastrulation as a potential origin of *Trim71*-KO related hematoendothelial phenotypes. At E7.5, *Trim71*-KO embryos are present at mendelian ratios (Torres-Fernández et al. 2021), appear normal in morphology and form a primitive streak. Mesodermal and endodermal cells were also present at similar numbers in E7.5 WT and *Trim71*-KO embryos, indicating that gastrulation occurs normally. The presence of EC in the yolk sac of *Trim71*-KO embryos further demonstrates that mesodermal progenitors are not impeded in their migration from the primitive streak to extraembryonic sites (Saykali et al. 2019). Nevertheless, our scRNA-seq data of E7.5 embryos show that widespread transcriptional differences are induced by *Trim71*-KO already at this developmental stage, preceding the appearance of morphological phenotypes by at least 1.5 days. This suggests that molecular changes in *Trim71*-KO embryos arise around gastrulation, providing a basis for the morphological phenotypes that become apparent at organogenesis. In the mesoderm, *Trim71*-KO led to decreased expression of genes involved in RNA splicing and chromatin remodeling. A previous study reported global changes in mRNA transcript splicing in *Trim71*-KO mESC, which was attributed to a direct Trim71-mediated repression of the splicing regulator Mbnl1 (Welte et al. 2019). While Mbnl1 was not identified as a DEG in the mesoderm in our scRNA-seq data, it is possible that Trim71 regulates RNA splicing during development by controlling the expression of multiple different splicing factors. Another recent study identified the regulation of the chromatin modifier cfp-1 by TRIM71 in *C. elegans* (Kumari et al. 2023), and the downregulation of chromatin remodeling factors in our data could hint at the control of epigenetic regulators as an additional conserved mechanism of how Trim71 shapes gene expression programs. Importantly, we show that loss of Trim71 leads to an increased expression of the transcription factor Eomes in the mesoderm *in vivo* and in differentiated mESC *in vitro*. We further identified a secondary structure in the 3ˈ UTR of the murine Eomes mRNA that fulfils the requirements of a TRE for the interaction with the NHL domain of Trim71 (Kumari et al. 2018). The location of this TRE in the Eomes mRNA is in line with the preferential positioning of TREs in the 3ˈ UTR of previously identified target mRNAs (Torres-Fernández et al. 2019; Welte et al. 2019; Kumari et al. 2023). The binding of Eomes mRNA by Trim71 was confirmed by CLIP and was abrogated by the R595H mutation in the RNA binding NHL domain of Trim71 (Duy et al. 2022), establishing Eomes as a direct target of Trim71-mediated repression in mesodermal development. In contrast, Lhx1 mRNA was devoid of putative TREs and was also not detectable after Trim71-FLAG-CLIP, demonstrating that Trim71 does not directly antagonize Lhx1 expression by mRNA binding. Transcription of the *Lhx1* locus is transactivated by Eomes (Nowotschin et al. 2013), thus the elevated Lhx1 mRNA levels in *Trim71*-KO embryos could be the result of the excessive Eomes expression in these embryos. Eomes is a crucial regulator of early embryonic development and loss of Eomes leads to defective epithelial-to-mesenchymal transition of nascent mesoderm, resulting in impaired mesoderm formation (Arnold et al. 2008; Russ et al. 2000). Moreover, Eomes is required for the generation of EryP from the mesoderm by influencing the chromatin landscape and enabling the accessibility of the DNA regions bound by the hematopoietic transcription factor SCL (Harland et al. 2021). The effects of excessive Eomes expression during mesodermal development *in vivo* are unknown, but it is reasonable to assume that accurate control of Eomes expression is vital for proper embryonic development. Thus, the direct regulation of Eomes mRNA levels shows a critical role of Trim71 in mesodermal development at gastrulation. In summary, we show that defects in primitive erythropoiesis and the cardiovascular system of *Trim71*-KO embryos are not caused by the loss of Trim71 in HEP, EC or EryP. Instead, our data strongly indicate that these phenotypes are caused by defects in the mesoderm during gastrulation that manifest at the onset of organogenesis.

## Materials and methods

### Mouse lines

All mice used in this study were bred in the licensed animal facility of the LIMES Institute (University of Bonn). The mouse lines *Trim71*-KO, *Trim71*-flox, *Csf1r*-iCre and *Tie2*-Cre used in this study have been described (Mitschka et al. 2015; Deng et al. 2010; Kisanuki et al. 2001). Conditional knockout of *Trim71* was induced by crossing female *Trim71*^fl/fl^ Cre-negative mice with male *Trim71*^fl/fl^ or *Trim71*^fl/+^ mice carrying a single copy of the Cre allele. Cre-positive *Trim71*^fl/fl^ animals were considered as the conditional knockout group, and Cre-negative littermates with either Trim71^fl/+^ or Trim71^fl/fl^ genotype were used as controls. Genotyping of mice and embryos was performed as described (Torres-Fernández et al. 2021). The ectoplacental cone was used for genotyping of E7.5–E8.5 embryos, while the embryo tail was used to genotype embryos from E9.5 on.

### Generation of mouse embryos

Mouse embryos were analyzed at developmental stages E7.5–E12.5 in this study. Timed mating was performed by placing one male and up to two female mice in a cage from afternoon until the morning of the next day. The presence of a vaginal plug was assessed as an indicator for mating, and a weight gain of more than 1.75 g of the female mouse was regarded as a sign of actual pregnancy (Galen W Heyne et al.). Pregnant female mice were sacrificed by cervical dislocation, and embryos were dissected in PBS under a SZX10 Stereomicroscope (Olympus) equipped with the cellSens Entry software (Olympus) for the acquisition of images.

### Analysis of the heartbeat in embryos

Videos of the embryo heartbeat were obtained during the preparation process with the cellSens Entry software (Olympus). Embryos that did not display any heart contractility for more than 60 s were classified as absent of a heartbeat. For embryos that showed heart contractility, at least three contractions were recorded and the heart rate was calculated as the mean duration between each contraction per minute.

### Flow cytometry and FACS of embryonic organs

Embryonic organs (yolk sac, embryo head, embryo body) were digested (100 mg/mL Collagenase D, 100 U/mL DNase I in 3% FCS/PBS) for 30 min at 37 °C, diluted with 2 mL FACS buffer (2 mM EDTA, 0.5% BSA in PBS) and minced through a 100 µm strainer to generate a single cell suspension (Iturri et al. 2017). Cells were centrifuged (320 g, 4 °C, 5 min), followed by incubation in Fc blocking solution (α-CD16/CD32 1:200 in FACS buffer) for 15 min at 4 °C and staining in antibody mix (all antibodies 1:200 in FACS buffer) for 30 min at 4 °C. Samples were washed and diluted 1:1 with DRAQ7 (1:1000 in FACS buffer) for dead cell exclusion before recording at the LSRII or FACS Symphony A5 flow cytometers (Becton Dickinson). Data analysis was performed in FlowJo (Becton Dickinson). Cell populations were identified by the combination of expressed surface markers: EryP (Ter119^+^ CD45^-^), EC (CD31^+^ AA4.1^-^), EMP (CD45^low^ Kit^+^ AA4.1^+^), pMac (CD45^+^ Kit^-^ CD11b^+^ F4/80^-^), macrophages (CD45^+^ Kit^-^ CD11b^+^ F4/80^+^).

Cell sorting was performed at an ARIA III (Becton Dickinson) with a 100 µm nozzle. 1000–5000 cells per sample were sorted into cooled 1.5 mL reaction tubes filled with 500 µl TRIzol Reagent (Thermo Fisher Scientific) and immediately transferred to -80 °C until further processing.

### Immunofluorescence staining and imaging

Embryonic organs were fixed in 4% PFA/PBS for 2h on a shaker at 4 °C and washed three times with PBS. For tissue sections, embryos were incubated with 30% sucrose in PBS for 48h, washed three times with PBS and embedded in O.C.T compound (Weckert Labortechnik) before freezing to -80 °C. 10–14 µm sections were prepared at a Cryostat CM3050S (Leica). Sections were dried at room temperature (rt) for 45 min, washed with PBS for 15 min and blocked for 1 h at rt in IF blocking buffer (5% normal goat serum, 0.3% Triton X-100, 0.5% BSA in PBS). Sections were stained with primary antibody mix (all antibodies 1:200 in PBS) overnight at 4 °C. Sections were then washed three times with PBS and incubated in secondary antibody mix (all antibodies 1:400 in PBS) including 1 ng/µl DAPI for 1 h at rt. After washing three times with PBS, sections were mounted with Fluoroshield (ImmunoBioScience). Yolk sacs were stained based on a published protocol (Roy and Delgado-Olguin 2018) as whole mounts floating in 48 well plates with one organ per well. The blocking and staining procedure for yolk sacs was the same as described for tissue sections, but incubation steps were performed on a shaker and the secondary antibody staining was extended to 90 min. Images were acquired using a LSM 880 Airyscan confocal microscope (Carl Zeiss). Image analysis was performed in ZEN blue (Carl Zeiss) and ImageJ. For the quantification of vascular structures in the yolk sac, three images per sample from different locations in the yolk sac were analyzed and the mean value was used as one data point. Endothelial extensions were defined as CD31^+^ structures that extend from an existing vessel without any connection to another vessel. Branching points were defined as the intersection of at least three vessels.

### Stem cell culture and differentiation

The *Trim71*-flox, *Trim71*-KO mESC lines and the mNeon-FLAG-tagged *Trim71*-flox, *Trim71*-KO and *Trim71*-R595H mESC lines have been described (Mitschka et al. 2015; Duy et al. 2022). Embryonic stem cells were maintained at 37 °C, 5% CO_2_ and 95% relative humidity on gelatin coated dishes in KnockOut DMEM (Thermo Fisher Scientific) supplemented with 15% FCS (Thermo Fisher Scientific), penicillin–streptomycin (Thermo Fisher Scientific), GlutaMAX (Thermo Fisher Scientific), MEM Non-Essential Amino Acids (Thermo Fisher Scientific), 50 µM β-Mercaptoethanol (Thermo Fisher Scientific), LIF (supernatant from L929 cells) and 2i CHIR99021 (3 µM, Sigma-Aldrich) and PD0325901 (1 µM, StemCell Technologies).

Differentiation of mESC was performed as embryoid bodies (EBs) in suspension by the removal of 2i and culturing in differentiation medium: StemPro-34 SFM (Thermo Fisher Scientific) supplemented with 10% FCS, penicillin–streptomycin, 2 mM L-Glutamine (PAN-Biotech), 40 µg/mL apo-Transferrin (Merck), 0.5 mM L-ascorbic acid (Sigma-Aldrich) and 0.15 mM 1-MTG (Sigma-Aldrich). To this end, adherent mESC were detached with Accutase (Thermo Fisher Scientific), washed twice with PBS and 2.5 x 10^6^ cells in 5 mL differentiation medium were seeded into a 60 mm non-treated petri dish (Greiner Bio-One). Cells were differentiated for 4 days with a medium change after 2 days without disruption of the EBs.

### Analysis of gene expression by qRT-PCR

RNA was extracted using TRIzol reagent (Thermo Fisher Scientific) according to manufacturer’s instructions. For low input samples, linear polyacrylamide (LPA) was added to the sample before RNA extraction (2.5 ng LPA for FACS-isolated cells, 7.5 ng LPA for CLIP samples). Isolated RNA was digested with 1 U/µl DNase I (Thermo Fisher Scientific) for 30 min at 37 °C, followed by addition of 1 µl 50 mM EDTA and heat inactivation for 10 min at 70 °C. 500 ng RNA were reverse transcribed using the High-Capacity cDNA Reverse Transcription Kit (Thermo Fisher Scientific) according to manufacturer’s instructions. Relative gene expression was quantified by qRT-PCR with TaqMan or SYBR Green assays (Bio-Rad) on a CFX96 thermal cycler (Bio-Rad). The following TaqMan probes were used: Trim71 (Mm01341471_m1), Eomes (Mm01351985_m1). For SYBR Green assays, the following primer pairs were used:

18S (for: GTAACCCGTTGAACCCCATTC, rev: CCATCCAATCGGTAGTAGCGAC)

Actb (for: CACTGTCGAGTCGCGTCC, rev: CGCAGCGATATCGTCATCCA)

Lhx1 (for: CGCCATATCCGTGAGCAACT, rev: CGCGCTTAGCTGTTTCATCC).

### Cross-linking RNA immunoprecipitation (CLIP)

CLIP was performed as described (Torres-Fernández et al. 2019). Briefly, for each condition the EBs from six 60 mm dishes of mESC expressing FLAG-tagged Trim71 variants at day 4 of differentiation were pooled, washed with cold PBS and UV-irradiated at 254 nm and 300 mJ/cm^2^. After centrifugation (320 g, 4 °C, 5 min), cells were lysed in TKM+ buffer for 15 min on ice (20 mM Tris, 100 mM KCl, 5 mM MgCl_2_, pH 7.4, supplemented with protease inhibitors 1:1000 PMSF, 1:1000 Benzamidin, 1:1000 Antipain; 1:2000 Aprotinin, 1:2000 Leupeptin, 0.2% NP-40 and RNase inhibitor 120 U/mL). Lysates were centrifuged (13200 rpm, 4 °C, 5 min) and a fraction of the supernatant was retained for protein and RNA input fractions. Equal amounts of protein from the supernatant (500–1500 mg) were immunoprecipitated with 30 µl anti-FLAG M2 magnetic beads (Sigma-Aldrich) at 4 °C on a spinning wheel for 4 h. After washing five times with TKM+ buffer, 20% of the IP fraction was used for western blot analysis, while 80% were digested with 0.5 mg/mL Proteinase K at 37 °C for 30 min and used for RNA extraction, cDNA generation and qRT-PCR analysis of target genes and 18S RNA as an unspecific binding reference gene. Enrichment values were calculated as enrichment = 2^ – [(Ct_CLIP_target_ – Ct_CLIP_ref_) – (Ct_Input_target_ – Ct_Input_ref_)] and normalized to the control genotype. Input and IP protein fractions were analyzed by western blot as described (Torres-Fernández et al. 2019). Briefly, protein lysates were separated by size via SDS-PAGE and transferred to a nitrocellulose membrane. Membranes were incubated over night at 4 °C with Trim71 (Worringer et al. 2014) or Vinculin (Sigma-Aldrich) antibodies (both 1:1000 in 5% milk powder in TBST: 50 mM Tris-HCl pH 7.6, 150 mM NaCl, 0.05% Tween-20), washed three times with TBST and incubated with species-matched HRP-coupled secondary antibodies (1:5000 in 5% milk powder in TBST) for 1 h at rt. After washing three times with TBST, membranes were developed with the Pierce ECL Substrate Kit (Thermo Fisher Scientific).

### Single-cell RNA-sequencing (scRNA-seq)

The 10x genomics Chromium Next GEM Single Cell 3ˈ Reagent Kits v3.1 (Dual Index) kit was used for scRNA-seq experiments. For developmental stage E7.5, one whole embryo of each genotype was used for sequencing. Single cells were isolated by digestion of E7.5 embryos with 0.25% Trypsin (Sigma-Aldrich) and 0.5 mM EDTA in PBS for 10 min at 37 °C, followed by mechanical dissociation by gentle resuspension through a 200 µl pipette tip and filtering through a 40 µm strainer into a 1.5 mL reaction tube. For E9.5 stage yolk sacs, cells isolated from two organs per genotype were pooled into one sample. Yolk sacs were digested with 100 mg/mL Collagenase D and 100 U/mL DNase I in 3% FCS/PBS for 30 min at 37 °C, diluted to 1.5 mL with PBS, minced through a 100 µm pore strainer and filtered through a 70 µm pore strainer into a 1.5 mL reaction tube. Cells isolated from E7.5 whole embryos or E9.5 yolk sacs in 1.5 mL reaction tubes were centrifuged (400g, 4 °C, 5 min) and completely loaded onto the Next GEM Chip G. Subsequent sample cleanup and library preparation was performed according to manufacturer’s instructions. Libraries were sequenced on a NovaSeq 6000 System (Illumina) with paired-end dual-indexing (28 cycles Read 1, 10 cycles i7, 10 cycles i5, 90 cycles Read 2) with NovaSeq 6000 S2 and SP (200 cycles) chemistry.

### Analysis of scRNA-seq data

Data generated in this study were processed using Cell Ranger v7.1.0 (10x Genomics). Specifically, raw sequencing data were demultiplexed with the cellranger mkfastq pipeline. The generated FASTQ files were further processed using the cellranger count pipeline for alignment, filtering, barcode counting, UMI counting and the generation of feature-barcode matrices. Mm10 2020A was used as a mouse reference genome. Subsequent data analysis was performed in R using Seurat (v.5.0) (Hao et al. 2024). Ambient RNA was removed by SoupX (Young and Behjati 2020) and high quality cells with 500–5000 features and less than 5% mitochondrial reads were filtered. The datasets of both genotypes were merged, normalized and scaled with standard settings from the Seurat package, followed by dimensionality reduction and visualization by UMAP. Cell types were annotated with SingleR and a reference dataset that was accessed via the MouseGastrulationData package (Pijuan-Sala et al. 2019). Cells from E7.5 embryos provided by this reference dataset were used to identify cell types in the E7.5 whole embryo scRNA-seq experiment. For the E9.5 yolk sac scRNA-seq experiment, cell type annotations from E8.5 embryos were used as a reference (Pijuan-Sala et al. 2019).

Hematopoietic cell populations in the yolk sac were further discriminated based on reported gene expression signatures for EMP, pMac and macrophages (Mass et al. 2016) as well as defined expression thresholds for *Maf* (macrophages) and *Pf4* (megakaryocytes). Differentially expressed genes between genotypes were identified for each cell population using the FindMarkers function from Seurat (Wilcoxon Rank Sum test), and filtered for an adjusted p-value of < 0.05 and an expression fold change of > 1.2 (upregulated) or < 0.8 (downregulated). Up- or downregulated DEG were used for gene ontology over representation analysis with the clusterProfiler package, accessing MSigDB gene annotations (Wu et al. 2021; Castanza et al. 2023). Expression scores for selected MSigDB processes were calculated with the AddModuleScore function from Seurat.

ScRNA-seq data of wildtype E6.5–E8.5 embryos was retrieved from the mouse gastrulation and early organogenesis cell atlas (Pijuan-Sala et al. 2019). Pseudo-bulk expression data was generated by accessing the data of embryonic stage E7.5 and E8.5 cells via the MouseGastrulationData package in R and using the AggregateExpression function from Seurat. Cells annotated as Erythroid3 in the source dataset were considered as EryP in the analysis.

### Prediction of RNA secondary structure

The sequence of the murine Eomes mRNA was downloaded from the NCBI (accession: NM_010136.4) and divided into 5ˈ UTR, coding sequence and 3ˈ UTR. Each region was further divided into 100 bp fragments that were individually analyzed with the RNAfold tool (http://rna.tbi.univie.ac.at/cgi-bin/RNAWebSuite/RNAfold.cgi) and displayed with color-coded positional entropy of each nucleotide.

## Data availability

The scRNA-seq data were submitted to the NCBI Gene Expression Omnibus with the accession number GSE272044.

## Acknowledgements

We thank Andrea Raths and Jordi Hees Soler for technical support. We are grateful to Shinya Yamanaka for providing the Trim71 antibody used in this study. Moreover, we are very thankful to Carmen Ruiz de Almodóvar for feedback on this manuscript. The work was funded by the Deutsche Forschungsgemeinschaft (DFG, German Research Foundation) under Germany’s Excellence Strategy-EXC2151-390873048 (to WK, EM, MB, AS), SFB 1454 – Project ID 432325352 (to WK, EM, AS, and MB), FOR5547 – Project-ID 503306912 (to EM) and the Bonner Forum Biomedizin (BFB Program for PhD students supporting innovative ideas, to TB). EM is supported by the European Research Council (ERC) under the European Union’s Horizon 2020 research and innovation program (Grant Agreement No. 851257). We would like to thank the Flow Cytometry Core Facility – Campus Poppelsdorf for providing support and instrumentation.

## Author contributions

TB and WK designed the study and experiments. TB performed the experiments and data analysis together with BJ (CLIP-experiments, *Csf1r^iCre^ Trim71* cKO embryos), HS (*Tie2^Cre^ Trim71* cKO embryos) and HT (scRNA-seq experiments). EDD, SP and MB performed sequencing of scRNA-seq cDNA libraries and pre-processed raw sequencing data. AS, EM and WK interpreted results and guided the experiments. TB wrote the manuscript with critical feedback from WK and all other authors. WK supervised the project.

## References

Arnold SJ, Hofmann UK, Bikoff EK, Robertson EJ. 2008: Pivotal roles for eomesodermin during axis formation, epithelium-to-mesenchyme transition and endoderm specification in the mouse. Development. 3: 501–511. doi: 10.1242/dev.014357.

Bardot ES, Hadjantonakis A-K. 2020: Mouse gastrulation: Coordination of tissue patterning, specification and diversification of cell fate. Mechanisms of Development 163: 103617. doi: 10.1016/j.mod.2020.103617.

Bautch VL, Caron KM. 2015: Blood and lymphatic vessel formation. Cold Spring Harbor Perspectives in Biology 3: a008268. doi: 10.1101/cshperspect.a008268.

Bazzoni G, Dejana E. 2004: Endothelial cell-to-cell junctions: molecular organization and role in vascular homeostasis. Physiological Reviews 3: 869–901. doi: 10.1152/physrev.00035.2003.

Bentley K, Franco CA, Philippides A, Blanco R, Dierkes M, Gebala V, Stanchi F, Jones M, Aspalter IM, Cagna G, et al. 2014: The role of differential VE-cadherin dynamics in cell rearrangement during angiogenesis. Nature Cell Biology 4: 309–321. doi: 10.1038/ncb2926.

Biben C, Weber TS, Potts KS, Choi J, Miles DC, Carmagnac A, Sargeant T, Graaf CA de, Fennell KA, Farley A, et al. 2023: In vivo clonal tracking reveals evidence of haemangioblast and haematomesoblast contribution to yolk sac haematopoiesis. Nature Communications 1: 41. doi: 10.1038/s41467-022-35744-x.

Castanza AS, Recla JM, Eby D, Thorvaldsdóttir H, Bult CJ, Mesirov JP. 2023: Extending support for mouse data in the Molecular Signatures Database (MSigDB). Nature Methods 11: 1619– 1620. doi: 10.1038/s41592-023-02014-7.

Chen J, Lai F, Niswander L. 2012: The ubiquitin ligase mLin41 temporally promotes neural progenitor cell maintenance through FGF signaling. Genes & Development 8: 803–815. doi: 10.1101/gad.187641.112.

Copp AJ, Greene NDE, Murdoch JN. 2003: The genetic basis of mammalian neurulation. Nature Reviews Genetics 10: 784–793. doi: 10.1038/nrg1181.

Coultas L, Chawengsaksophak K, Rossant J. 2005: Endothelial cells and VEGF in vascular development. Nature 7070: 937–945. doi: 10.1038/nature04479.

Cuevas E, Rybak-Wolf A, Rohde AM, Nguyen DTT, Wulczyn FG. 2015: Lin41/Trim71 is essential for mouse development and specifically expressed in postnatal ependymal cells of the brain. Frontiers in Cell and Developmental Biology 3: 20. doi: 10.3389/fcell.2015.00020.

Deng L, Zhou J-F, Sellers RS, Li J-F, Nguyen AV, Wang Y, Orlofsky A, Liu Q, Hume DA, Pollard JW, et al. 2010: A novel mouse model of inflammatory bowel disease links mammalian target of rapamycin-dependent hyperproliferation of colonic epithelium to inflammation-associated tumorigenesis. The American Journal of Pathology 2: 952–967. doi: 10.2353/ajpath.2010.090622.

Du G, Wang X, Luo M, Xu W, Zhou T, Wang M, Yu L, Li L, Cai L’e, Wang PJ, et al. 2020: mRBPome capture identifies the RNA-binding protein TRIM71, an essential regulator of spermatogonial differentiation. Development. 8: doi: 10.1242/dev.184655.

Dumont DJ, Gradwohl G, Fong GH, Puri MC, Gertsenstein M, Auerbach A, Breitman ML. 1994: Dominant-negative and targeted null mutations in the endothelial receptor tyrosine kinase, tek, reveal a critical role in vasculogenesis of the embryo. Genes & Development 16: 1897– 1909. doi: 10.1101/gad.8.16.1897.

Duy PQ, Jux B, Zhao S, Mekbib KY, Dennis E, Dong W, Nelson-Williams C, Mehta NH, Shohfi JP, Juusola J, et al. 2024: TRIM71 mutations cause a neurodevelopmental syndrome featuring ventriculomegaly and hydrocephalus. Brain epub ahead of print: doi: 10.1093/brain/awae175.

Duy PQ, Weise SC, Marini C, Li X-J, Liang D, Dahl PJ, Ma S, Spajic A, Dong W, Juusola J, et al. 2022: Impaired neurogenesis alters brain biomechanics in a neuroprogenitor-based genetic subtype of congenital hydrocephalus. Nature Neuroscience 4: 458–473. doi: 10.1038/s41593-022-01043-3.

Ecsedi M, Grosshans H. 2013: LIN-41/TRIM71: emancipation of a miRNA target. Genes & Development 6: 581–589. doi: 10.1101/gad.207266.112.

Ecsedi M, Rausch M, Großhans H. 2015: The let-7 microRNA directs vulval development through a single target. Developmental Cell 3: 335–344. doi: 10.1016/j.devcel.2014.12.018.

Ema M, Yokomizo T, Wakamatsu A, Terunuma T, Yamamoto M, Takahashi S. 2006: Primitive erythropoiesis from mesodermal precursors expressing VE-cadherin, PECAM-1, Tie2, endoglin, and CD34 in the mouse embryo. Blood 13: 4018–4024. doi: 10.1182/blood-2006-03-012872.

Fujiwara Y, Browne CP, Cunniff K, Goff SC, Orkin SH. 1996: Arrested development of embryonic red cell precursors in mouse embryos lacking transcription factor GATA-1. Proceedings of the National Academy of Sciences of the United States of America 22: 12355–12358. doi: 10.1073/pnas.93.22.12355.

Galen W Heyne, Erin H Plisch, Cal G Melberg, Eric P Sandgren, Jody A Peter, Robert J Lipinski: A Simple and Reliable Method for Early Pregnancy Detection in Inbred Mice:

Garcia MD, Larina IV. 2014: Vascular development and hemodynamic force in the mouse yolk sac. Frontiers in Physiology 5: 308. doi: 10.3389/fphys.2014.00308.

Gavrilov S, Lacy E. 2013: Genetic dissection of ventral folding morphogenesis in mouse: embryonic visceral endoderm-supplied BMP2 positions head and heart. Current Opinion in Genetics & Development 4: 461–469. doi: 10.1016/j.gde.2013.04.001.

Ginhoux F, Greter M, Leboeuf M, Nandi S, See P, Gokhan S, Mehler MF, Conway SJ, Ng LG, Stanley ER, et al. 2010: Fate Mapping Analysis Reveals That Adult Microglia Derive from Primitive Macrophages. Science 6005: 841–845. doi: 10.1126/science.1194637.

Hao Y, Stuart T, Kowalski MH, Choudhary S, Hoffman P, Hartman A, Srivastava A, Molla G, Madad S, Fernandez-Granda C, et al. 2024: Dictionary learning for integrative, multimodal and scalable single-cell analysis. Nature Biotechnology 2: 293–304. doi: 10.1038/s41587-023-01767-y.

Harland LTG, Simon CS, Senft AD, Costello I, Greder L, Imaz-Rosshandler I, Göttgens B, Marioni JC, Bikoff EK, Porcher C, et al. 2021: The T-box transcription factor Eomesodermin governs haemogenic competence of yolk sac mesodermal progenitors. Nature Cell Biology 1: 61–74. doi: 10.1038/s41556-020-00611-8.

Iturri L, Freyer L, Biton A, Dardenne P, Lallemand Y, Gomez Perdiguero E. 2021: Megakaryocyte production is sustained by direct differentiation from erythromyeloid progenitors in the yolk sac until midgestation. Immunity 7: 1433–1446.e5. doi: 10.1016/j.immuni.2021.04.026.

Iturri L, Saenz Coronilla J, Lallemand Y, Gomez Perdiguero E. 2017: Identification Of Erythromyeloid Progenitors And Their Progeny In The Mouse Embryo By Flow Cytometry. Journal of visualized experiments : JoVE 125: doi: 10.3791/55305.

Jones EAV, Yuan L, Breant C, Watts RJ, Eichmann A. 2008: Separating genetic and hemodynamic defects in neuropilin 1 knockout embryos. Development. 14: 2479–2488. doi: 10.1242/dev.014902.

Kasaai B, Caolo V, Peacock HM, Lehoux S, Gomez-Perdiguero E, Luttun A, Jones EAV. 2017: Erythro-myeloid progenitors can differentiate from endothelial cells and modulate embryonic vascular remodeling. Scientific Reports 7: 43817. doi: 10.1038/srep43817.

Kisanuki YY, Hammer RE, Miyazaki J, Williams SC, Richardson JA, Yanagisawa M. 2001: Tie2-Cre transgenic mice: a new model for endothelial cell-lineage analysis in vivo. Developmental Biology 2: 230–242. doi: 10.1006/dbio.2000.0106.

Koushik SV, Wang J, Rogers R, Moskophidis D, Lambert NA, Creazzo TL, Conway SJ. 2001: Targeted inactivation of the sodium-calcium exchanger (Ncx1) results in the lack of a heartbeat and abnormal myofibrillar organization. FASEB journal: Official Publication of the Federation of American Societies for Experimental Biology 7: 1209–1211. doi: 10.1096/fj.00-0696fje.

Krebs LT, Norton CR, Shutter JR, Maguire M, Sundberg JP, Gallahan D, Closson V, Kitajewski J, Callahan R, Smith GH, et al. 2000: Notch signaling is essential for vascular morphogenesis in mice. Genes & Development 14: 1343–1352. doi: 10.1101/gad.14.11.1343.

Kumari P, Aeschimann F, Gaidatzis D, Keusch JJ, Ghosh P, Neagu A, Pachulska-Wieczorek K, Bujnicki JM, Gut H, Großhans H, et al. 2018: Evolutionary plasticity of the NHL domain underlies distinct solutions to RNA recognition. Nature Communications 1: 1549. doi: 10.1038/s41467-018-03920-7.

Kumari P, Thuestad LH, Ciosk R. 2023: Post-transcriptional repression of CFP-1 expands the regulatory repertoire of LIN-41/TRIM71. Nucleic Acids Research 19: 10668–10680. doi: 10.1093/nar/gkad729.

Lamalice L, Le Boeuf F, Huot J. 2007: Endothelial cell migration during angiogenesis. Circulation Research 6: 782–794. doi: 10.1161/01.RES.0000259593.07661.1e.

Lee JS, Yu Q, Shin JT, Sebzda E, Bertozzi C, Chen M, Mericko P, Stadtfeld M, Zhou D, Cheng L, et al. 2006: Klf2 is an essential regulator of vascular hemodynamic forces in vivo. Developmental Cell 6: 845–857. doi: 10.1016/j.devcel.2006.09.006.

Lelièvre E, Mattot V, Huber P, Vandenbunder B, Soncin F. 2000: ETS1 lowers capillary endothelial cell density at confluence and induces the expression of VE-cadherin. Oncogene 20: 2438–2446. doi: 10.1038/sj.onc.1203563.

Lin Y-C, Hsieh L-C, Kuo M-W, Yu J, Kuo H-H, Lo W-L, Lin R-J, Yu AL, Li W-H. 2007: Human TRIM71 and its nematode homologue are targets of let-7 microRNA and its zebrafish orthologue is essential for development. Molecular Biology and Evolution 11: 2525–2534. doi: 10.1093/molbev/msm195.

Loedige I, Gaidatzis D, Sack R, Meister G, Filipowicz W. 2013: The mammalian TRIM-NHL protein TRIM71/LIN-41 is a repressor of mRNA function. Nucleic Acids Research 1: 518–532. doi: 10.1093/nar/gks1032.

Lux CT, Yoshimoto M, McGrath K, Conway SJ, Palis J, Yoder MC. 2008: All primitive and definitive hematopoietic progenitor cells emerging before E10 in the mouse embryo are products of the yolk sac. Blood 7: 3435–3438. doi: 10.1182/blood-2007-08-107086.

Maller Schulman BR, Liang X, Stahlhut C, DelConte C, Stefani G, Slack FJ. 2008: The let-7 microRNA target gene, Mlin41/Trim71 is required for mouse embryonic survival and neural tube closure. Cell Cycle 24: 3935–3942. doi: 10.4161/cc.7.24.7397.

Mass E, Ballesteros I, Farlik M, Halbritter F, Günther P, Crozet L, Jacome-Galarza CE, Händler K, Klughammer J, Kobayashi Y, et al. 2016: Specification of tissue-resident macrophages during organogenesis. Science 6304: doi: 10.1126/science.aaf4238.

McGrath K, Palis J. 2008: Ontogeny of Erythropoiesis in the Mammalian Embryo. Current Topics in Developmental Biology 82: 1–22. doi: 10.1016/S0070-2153(07)00001-4.

McGrath KE, Frame JM, Fegan KH, Bowen JR, Conway SJ, Catherman SC, Kingsley PD, Koniski AD, Palis J. 2015: Distinct Sources of Hematopoietic Progenitors Emerge before HSCs and Provide Functional Blood Cells in the Mammalian Embryo. Cell Reports 12: 1892–1904. doi: 10.1016/j.celrep.2015.05.036.

McGrath KE, Koniski AD, Malik J, Palis J. 2003: Circulation is established in a stepwise pattern in the mammalian embryo. Blood 5: 1669–1676. doi: 10.1182/blood-2002-08-2531.

Mitschka S, Ulas T, Goller T, Schneider K, Egert A, Mertens J, Brüstle O, Schorle H, Beyer M, Klee K, et al. 2015: Co-existence of intact stemness and priming of neural differentiation programs in mES cells lacking Trim71. Scientific Reports 5: 11126. doi: 10.1038/srep11126.

Nowotschin S, Costello I, Piliszek A, Kwon GS, Mao C-a, Klein WH, Robertson EJ, Hadjantonakis A-K. 2013: The T-box transcription factor Eomesodermin is essential for AVE induction in the mouse embryo. Genes & Development 9: 997–1002. doi: 10.1101/gad.215152.113.

Pearson S, Sroczynska P, Lacaud G, Kouskoff V. 2008: The stepwise specification of embryonic stem cells to hematopoietic fate is driven by sequential exposure to Bmp4, activin A, bFGF and VEGF. Development. 8: 1525–1535. doi: 10.1242/dev.011767.

Phoon CK, Aristizabal O, Turnbull DH. 2000: 40 MHz Doppler characterization of umbilical and dorsal aortic blood flow in the early mouse embryo. Ultrasound in Medicine and Biology 8: 1275–1283. doi: 10.1016/S0301-5629(00)00278-7.

Pijuan-Sala B, Griffiths JA, Guibentif C, Hiscock TW, Jawaid W, Calero-Nieto FJ, Mulas C, Ibarra-Soria X, Tyser RCV, Ho DLL, et al. 2019: A single-cell molecular map of mouse gastrulation and early organogenesis. Nature 7745: 490–495. doi: 10.1038/s41586-019-0933-9.

Prummel KD, Nieuwenhuize S, Mosimann C. 2020: The lateral plate mesoderm. Development. 12: doi: 10.1242/dev.175059.

Roy AR, Delgado-Olguin P. 2018: Visualizing the Vascular Network in the Mouse Embryo and Yolk Sac. Methods in Molecular Biology 1752: 11–16. doi: 10.1007/978-1-4939-7714-7_2.

Russ AP, Wattler S, Colledge WH, Aparicio SA, Carlton MB, Pearce JJ, Barton SC, Surani MA, Ryan K, Nehls MC, et al. 2000: Eomesodermin is required for mouse trophoblast development and mesoderm formation. Nature 6773: 95–99. doi: 10.1038/35003601.

Saykali B, Mathiah N, Nahaboo W, Racu M-L, Hammou L, Defrance M, Migeotte I. 2019: Distinct mesoderm migration phenotypes in extra-embryonic and embryonic regions of the early mouse embryo. eLife 8: doi: 10.7554/eLife.42434.

Shalaby F, Rossant J, Yamaguchi TP, Gertsenstein M, Wu XF, Breitman ML, Schuh AC. 1995: Failure of blood-island formation and vasculogenesis in Flk-1-deficient mice. Nature 6535: 62–66. doi: 10.1038/376062a0.

Shi F, Zhang K, Cheng Q, Che S, Zhi S, Yu Z, Liu F, Duan F, Wang Y, Yang N. 2024: Molecular mechanism governing RNA-binding property of mammalian TRIM71 protein. Science Bulletin 1: 72–81. doi: 10.1016/j.scib.2023.11.041.

Stremmel C, Schuchert R, Wagner F, Thaler R, Weinberger T, Pick R, Mass E, Ishikawa-Ankerhold HC, Margraf A, Hutter S, et al. 2018: Yolk sac macrophage progenitors traffic to the embryo during defined stages of development. Nature Communications 1: 75. doi: 10.1038/s41467-017-02492-2.

Tachibana K, Jones N, Dumont DJ, Puri MC, Bernstein A. 2005: Selective role of a distinct tyrosine residue on Tie2 in heart development and early hematopoiesis. Molecular and Cellular Biology 11: 4693–4702. doi: 10.1128/MCB.25.11.4693-4702.2005.

Tang Y, Harrington A, Yang X, Friesel RE, Liaw L. 2010: The contribution of the Tie2+ lineage to primitive and definitive hematopoietic cells. Genesis 9: 563–567. doi: 10.1002/dvg.20654.

Torres-Fernández LA, Emich J, Port Y, Mitschka S, Wöste M, Schneider S, Fietz D, Oud MS, Di Persio S, Neuhaus N, et al. 2021: TRIM71 Deficiency Causes Germ Cell Loss During Mouse Embryogenesis and Is Associated With Human Male Infertility. Frontiers in Cell and Developmental Biology 9: 658966. doi: 10.3389/fcell.2021.658966.

Torres-Fernández LA, Jux B, Bille M, Port Y, Schneider K, Geyer M, Mayer G, Kolanus W. 2019: The mRNA repressor TRIM71 cooperates with Nonsense-Mediated Decay factors to destabilize the mRNA of CDKN1A/p21. Nucleic Acids Research 22: 11861–11879. doi: 10.1093/nar/gkz1057.

Völkers M, Preiss T, Hentze MW. 2024: RNA-binding proteins in cardiovascular biology and disease: the beat goes on. Nature Reviews Cardiology 21: doi: 10.1038/s41569-023-00958-z.

Wei G, Srinivasan R, Cantemir-Stone CZ, Sharma SM, Santhanam R, Weinstein M, Muthusamy N, Man AK, Oshima RG, Leone G, et al. 2009: Ets1 and Ets2 are required for endothelial cell survival during embryonic angiogenesis. Blood 5: 1123–1130. doi: 10.1182/blood-2009-03-211391.

Welte T, Goulois A, Stadler MB, Hess D, Soneson C, Neagu A, Azzi C, Wisser MJ, Seebacher J, Schmidt I, et al. 2023: Convergence of multiple RNA-silencing pathways on GW182/TNRC6. Molecular Cell 14: 2478–2492.e8. doi: 10.1016/j.molcel.2023.06.001.

Welte T, Tuck AC, Papasaikas P, Carl SH, Flemr M, Knuckles P, Rankova A, Bühler M, Großhans H. 2019: The RNA hairpin binder TRIM71 modulates alternative splicing by repressing MBNL1. Genes & Development 33: 1221–1235. doi: 10.1101/gad.328492.119.

Worringer KA, Rand TA, Hayashi Y, Sami S, Takahashi K, Tanabe K, Narita M, Srivastava D, Yamanaka S. 2014: The let-7/LIN-41 pathway regulates reprogramming to human induced pluripotent stem cells by controlling expression of prodifferentiation genes. Cell Stem Cell 1: 40–52. doi: 10.1016/j.stem.2013.11.001.

Wu T, Hu E, Xu S, Chen M, Guo P, Dai Z, Feng T, Zhou L, Tang W, Zhan L, et al. 2021: clusterProfiler 4.0: A universal enrichment tool for interpreting omics data. The Innovation 3: 100141. doi: 10.1016/j.xinn.2021.100141.

Young MD, Behjati S. 2020: SoupX removes ambient RNA contamination from droplet-based single-cell RNA sequencing data. GigaScience 12: doi: 10.1093/gigascience/giaa151.

