## Supplement for "Dysregulation of gene expression during gastrulation results in impaired primitive erythropoiesis and vascular development in Trim71-KO embryos"

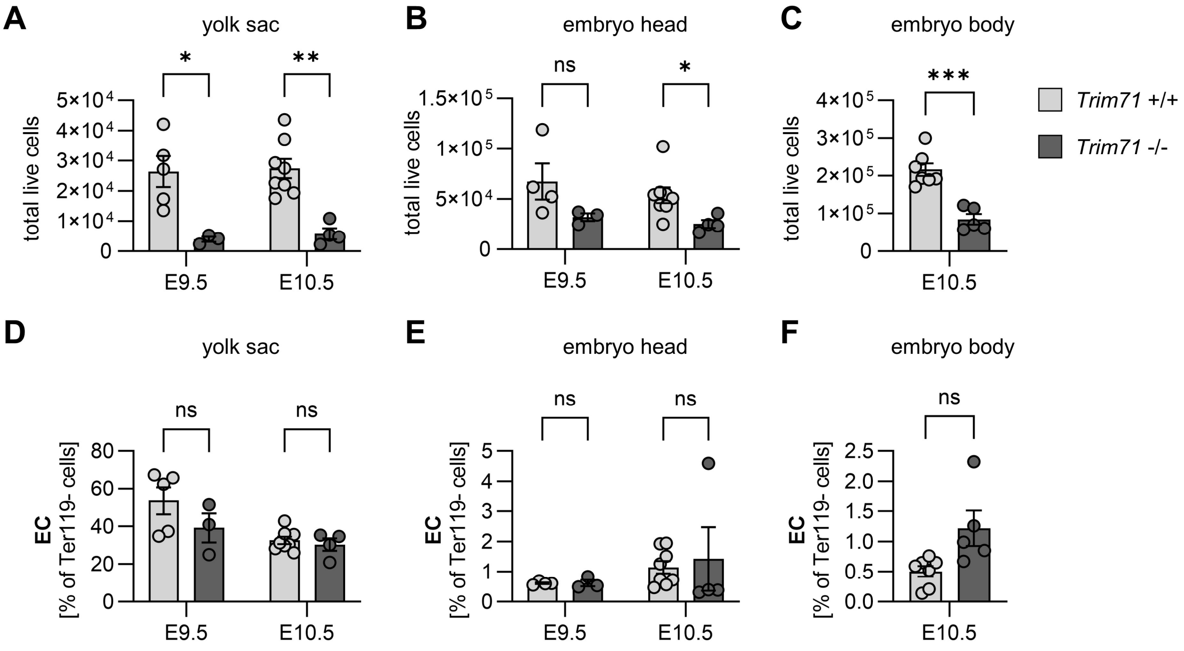


Supplemental Figure 1: Effect of *Trim71*-KO on total live cell numbers and relative EC numbers. (A–C) Flow cytometric quantification of total live cells in the (A) yolk sac and (B) embryo head at E9.5 and (C) the embryo body at E10.5. (D–F) Flow cytometric quantification of relative EC numbers in the (D) yolk sac and (E) embryo head at E9.5 and E10.5 and (F) the embryo body at E10.5 (n = 3–8 embryos from 2–3 experiments, data depicted as mean±SEM, unpaired Student's *t*-test, ns = not significant, (*) $P < 0.05$, (**) $P < 0.01$, (***) $P < 0.001$).

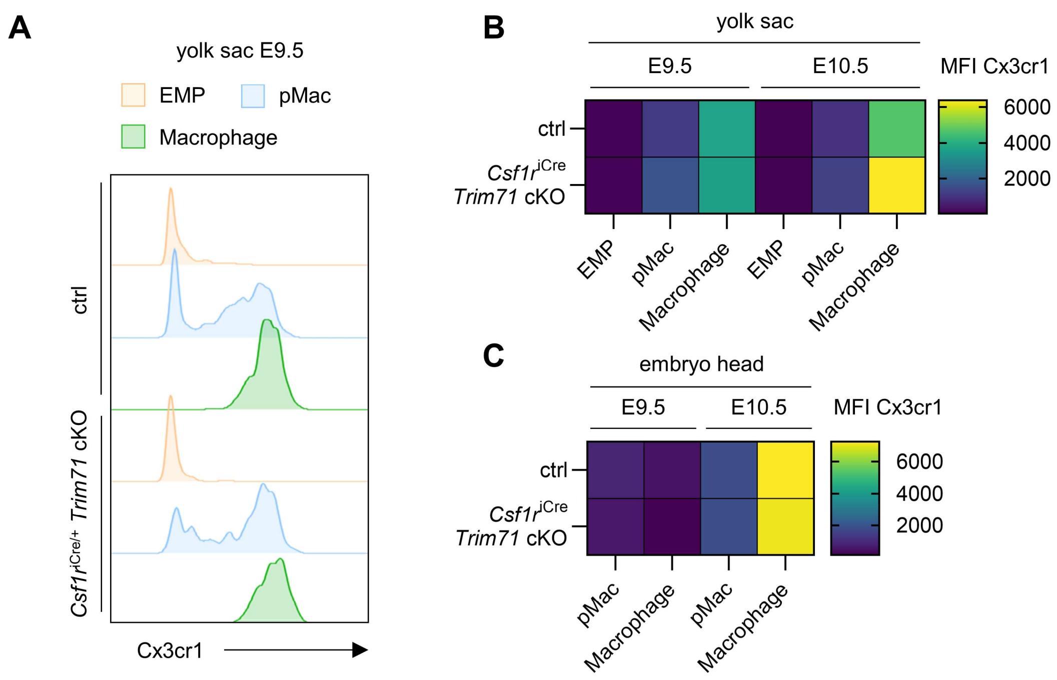


Supplemental Figure 2: *Csf1r*$^{iCre}$ *Trim71* cKO does not influence Cx3cr1 expression in pMac and macrophages. (A) Representative flow cytometry histograms of Cx3cr1 fluorescence intensity in E9.5 yolk sac EMP, pMac and macrophages. (B, C) MFI of Cx3cr1 in (B) EMP, pMac and macrophages of the yolk sac at E9.5 and E10.5 and (C) pMac and macrophages of the embryo head at E9.5 and E10.5 (n = 9–15 embryos from 4–5 experiments, data depicted as geometric mean).

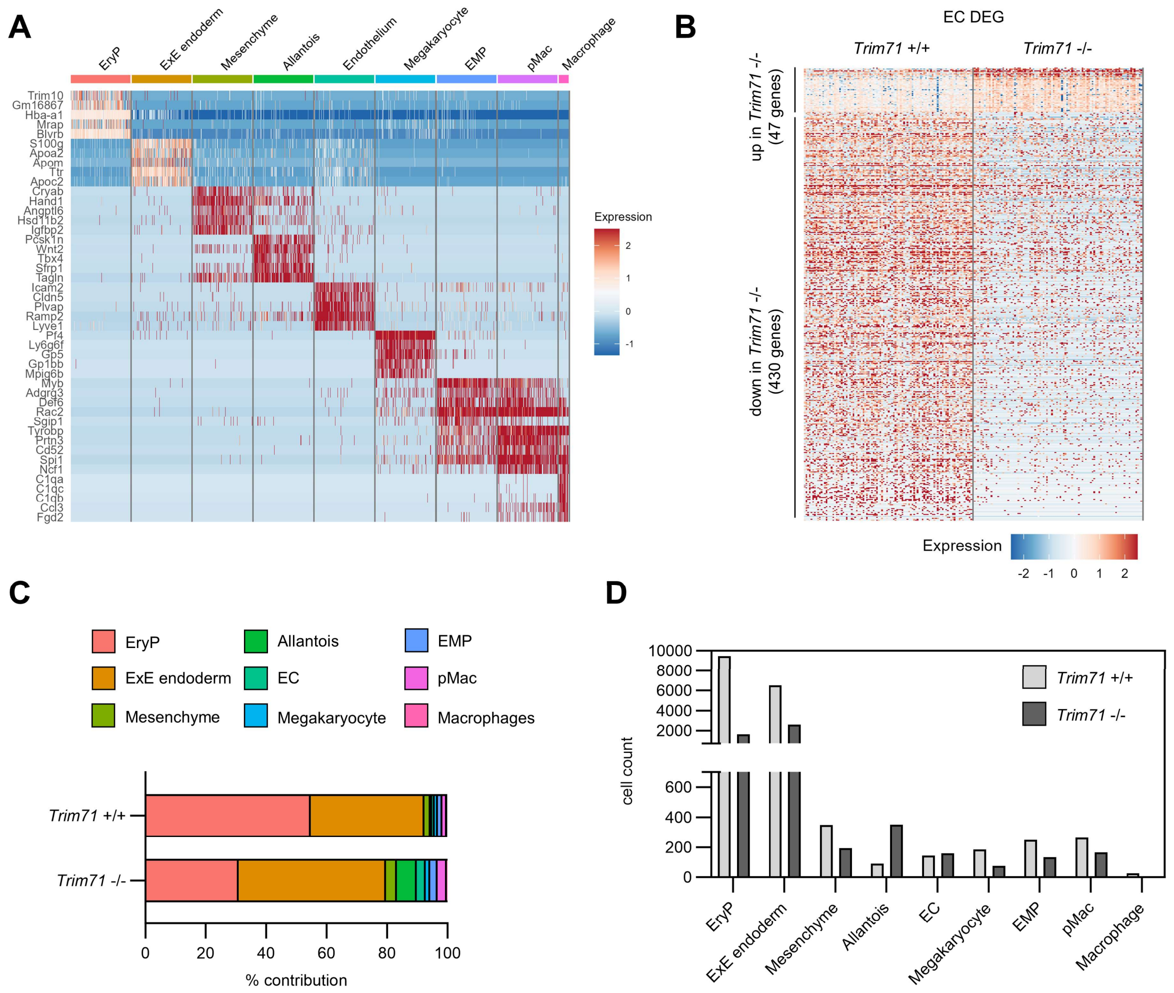


Supplemental Figure 3: Cell type marker genes and cell numbers as determined by scRNA-seq of E9.5 yolk sacs. (A) Expression heatmap of top marker genes across all cell types from scRNA-seq of *Trim71*-WT or *Trim71*-KO E9.5 yolk sacs. (B) Expression heatmap of upregulated and downregulated DEG in *Trim71*$^{+/+}$and *Trim71*$^{-/-}$ EC. (C) Percentage contribution of cell types to total cells by genotype. (D) Absolute cell counts of each cell type by genotype.

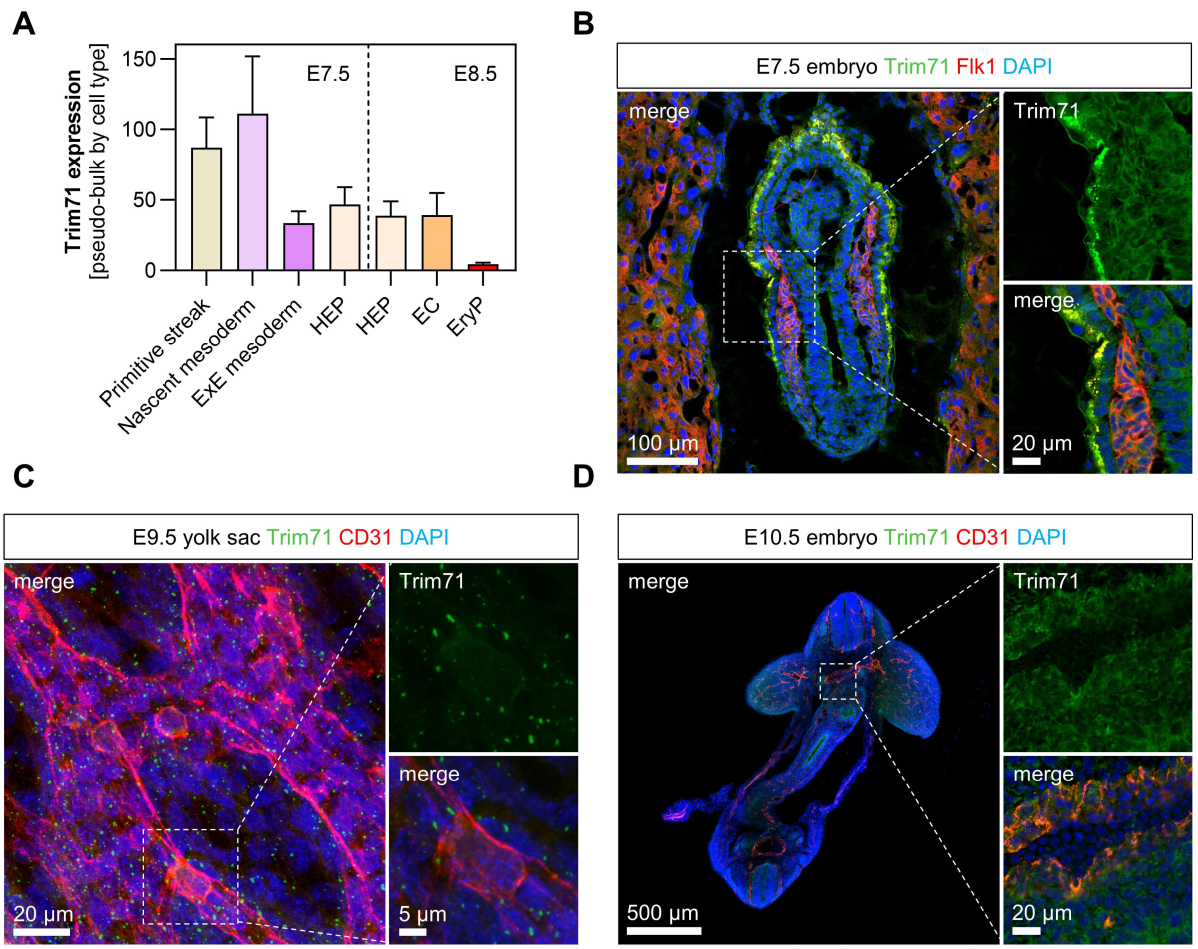


Supplemental Figure 4: Expression of Trim71 at gastrulation and early organogenesis in the hematoendothelial lineage. (A) Analysis of data from the WT E6.5–E8.5 mouse cell atlas from Pijuan-Sala et al. (2019). Trim71 expression in selected cell types from E7.5 or E8.5 embryos after pseudo-bulk aggregation of gene expression data by cell types from biological replicates (n = 3–4). (B) Immunofluorescence staining of Trim71 (green), Flk1 (red) and DAPI (blue) in a sagittal section of an E7.5 WT embryo. Dashed box shows magnification of Flk1$^{+}$ cells. (C) Immunofluorescence staining of Trim71 (green), CD31 (red) and DAPI (blue) in an E9.5 WT yolk sac. Dashed box shows magnification of a CD31$^{+}$ EC. (D) Immunofluorescence staining of Trim71 (green), CD31 (red) and DAPI (blue) in a transversal section of an E10.5 WT embryo. Dashed box shows magnification of the dorsal aorta.

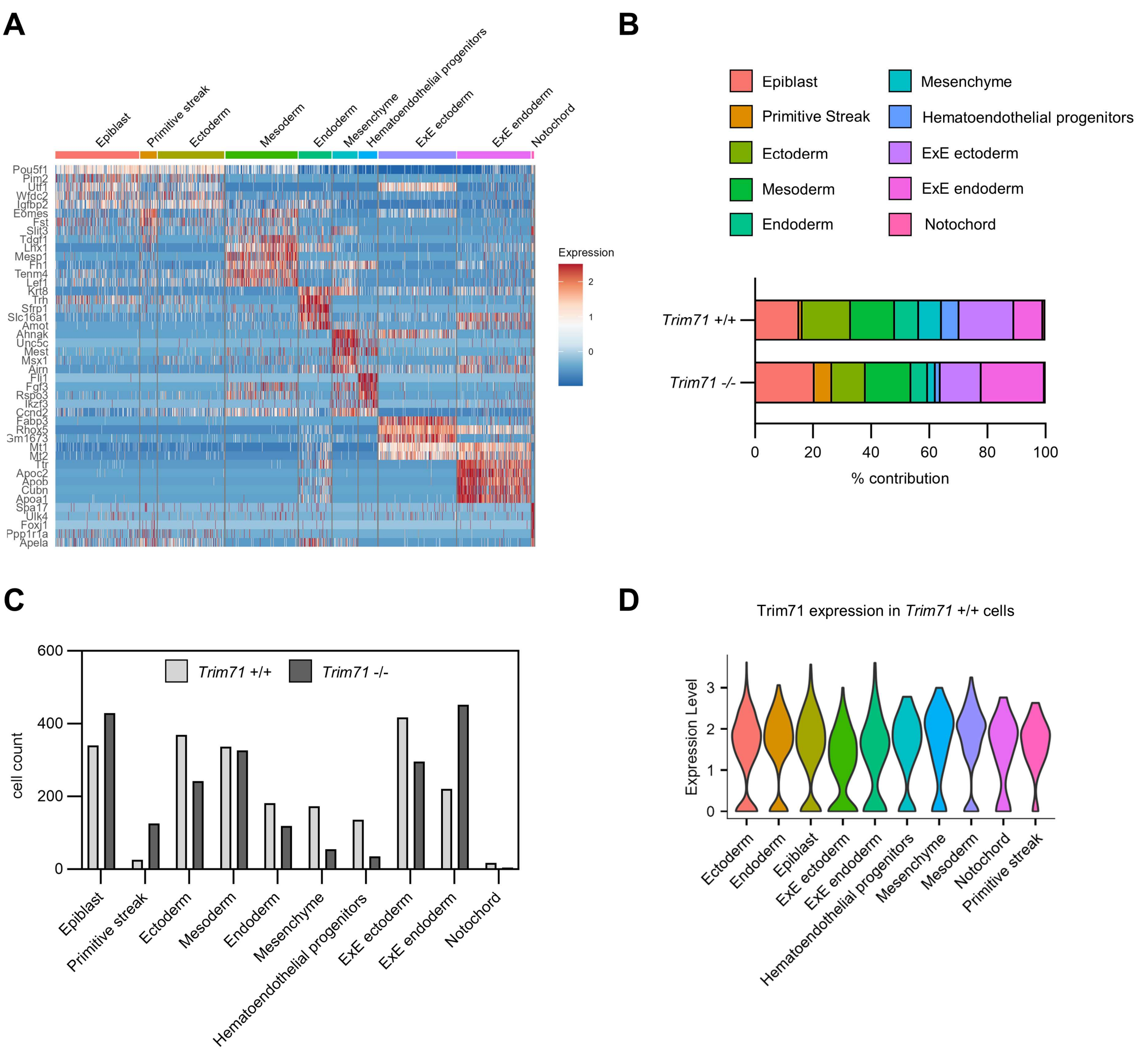


Supplemental Figure 5: Cell type marker genes and cell numbers as determined by scRNA-seq of E7.5 whole embryos. (A) Expression heatmap of top marker genes across all cell types from scRNA-seq of *Trim71*-WT or *Trim71*-KO E7.5 whole embryos. (B) Percentage contribution of cell types to total cells by genotype. (C) Absolute cell counts of each cell type by genotype. (D) Expression of Trim71 in all cell types in *Trim71*$^{+/+}$ cells.

Supplemental Video 1: *Trim71*-WT E10.5 embryo with present heartbeat and normal heart rate.

Supplemental Video 2: *Trim71*-KO E10.5 embryo with absent heartbeat.

Supplemental Video 3: *Trim71*-KO E10.5 embryo with present heartbeat but decreased heart rate.
